# How Aberrant *N*-Glycosylation Can Alter Protein Functionality and Ligand Binding: an Atomistic View

**DOI:** 10.1101/2022.12.22.521543

**Authors:** Matteo Castelli, Pengrong Yan, Anna Rodina, Chander S. Digwal, Palak Panchal, Gabriela Chiosis, Elisabetta Moroni, Giorgio Colombo

**Author notes:** Correspondence (G.Colombo.); (E.M.); (G.Chiosis.).

## Abstract

Protein assembly defects due to enrichment of aberrant conformational variants of proteins are emerging as a new frontier in therapeutics design. Understanding, atomistically, structural elements that remodel the energy landscape of proteins, with the consequence of rewiring the conformational dynamics of proteins and pathologically perturbing functionally-oriented ensembles, is key for development of inhibitors. This is particularly relevant for molecular chaperones, hub proteins for the assembly of large multiprotein complexes, where enrichment of aberrant conformers can have a large impact on the cellular proteome, and in turn, on phenotypes. Here, we integrate computational and experimental tools to unveil how *N*-glycosylation of specific residues in glucose-regulated protein 94 (GRP94) modulates internal dynamics and alters the conformational fitness of regions fundamental for interaction with the nucleotide and synthetic ligands, and impacts substructures dedicated to recognition of interacting proteins. We show how *N*-glycosylation plays an active role in modulating the energy landscape of the protein, with specific glycosylation patterns determining specific functionally-oriented dynamic signatures. Our results provide support for leveraging the structural-dynamics knowledge on distinct glycosylation variants to design molecules targeting GRP94 disease-associated conformational states and assemblies. Since glycosylation is the most abundant form of post-translational modification, our results and mechanistic models can readily be transferred to other targets and contexts for cancers and other diseases.

## Introduction

Protein function, and in turn malfunction, is determined by the extent conformational ensembles populate distinct states, each endowed with specific activities, and by the mechanisms of dynamic transition from a conformational state to another^1–7^. Changes in the distribution of such conformational states is governed by both non-covalent events, such as allosteric ligand binding, and covalent events such as post-translational modifications (PTMs)^8,9^. These modifications have evolved as a mechanism of adaptation to select specific functionally-oriented protein ensembles, either by stabilizing or destabilizing active and inactive conformations.

The importance of PTMs cannot be overstated, and recent investigations have started to shed light on their significance in regulating both physiological and disease conditions^10–12^. In the context of a dynamic model of regulation of protein conformational ensembles, PTMs work by shifting populations of protein ensembles from a functional state to another, and thus modulate the abundance of specific forms. This may have a dramatic impact on cellular networks, as different conformers may engage in distinct interactions in the context of multiprotein complexes, in turn remodeling protein-protein interactions (PPI) and the assembly of the large multimolecular assemblies required to regulate biomolecular pathways.

Among PTMs, glycosylation plays a host of important roles^13–15^. This fact is vividly illustrated by recent studies on the Sars-CoV-2 Spike protein, whereby glycosylation of specific positions, and the types of carbohydrate chains attached, not only function to shield the protein from the host immune system, but also play an active part in modulating the stability of the protein and the accessibility of the receptor binding domain, mechanisms which are at the basis of viral entry^16,17^. Glycosylation has also emerged as a mechanism in cancers and inflammation where cellular localization as well as protein conformations and assemblies could be pathologically remodeled by this PTM^10,15,18^. For example, we recently reported on a disease-specific variant of the chaperone GRP94, where, by altering *N*-glycosylation, a new protein that is conformationally, dynamically, and functionally distinct from the GRP94 of normal cells is created^15^. In place of a chaperone which is unglycosylated or monoglycosylated on N217, is confined to the ER and folds client proteins through transient interactions, a specific increase of *N*-glycosylation on N62 promotes a conformational state in GRP94 that allows for plasma membrane (PM) localization and stable interactions with PM proteins^15^. Through this complexation stabilization, the functions of PM proteins are enhanced, and downstream cellular protein pathways are aberrantly remodeled. *N*-glycosylation thus transforms a chaperone, GRP94, from a folding to a scaffolding protein that in turn remodels protein assembly and connectivity, with an end result of proteome-wide dysfunction^19,20^.

Aggregating these findings, aberrantly glycosylated protein variants and their altered assemblies have emerged as promising pharmacological targets in infectious diseases, inflammation and cancer^15,19–23^. Despite such growing evidence on the importance of glycosylation as a disease mechanism, an atomistic view on how glycosylation may influence proteins is largely missing. Specifically, the molecular mechanisms underlying the modulation of conformational dynamics and of molecular recognition properties with regards to substrate and drug-like inhibitors, as well as their impact on functions remain poorly understood. Without this knowledge, the development of inhibitors and clinical candidates targeting specific disease-associated, glycosylation-dependent, conformational states and assemblies is hampered.

Here, we set out to unveil at atomistic resolution the mechanism through which *N*-glycans at different protein sites modulate the conformational dynamics of GRP94, affecting both its interaction mode with clients and nucleotide processing. The degree of glycosylation and the specific positioning of the glycosyl moieties are thus reconnected to their impact on function. On this basis, we rationalize the effects of distinct ligands on different forms of GRP94. In this context, we document the reach and implications of our approach through chemical biology tests of the activity of different designed molecules. Specifically, we use long-timescale Molecular Dynamics (MD) simulations (around 40 micro-seconds in total) to expose the links between protein dynamics and the establishment of specific interaction patterns in different protein forms, and leverage this knowledge to build a simple model of PTM-induced functional perturbation mechanisms. Finally, we use these results to rationalize productive and nonproductive engagement of conformational states by small molecule ligands, thus providing key insights into the design of inhibitors targeting the disease-specific, aberrantly glycosylated, GRP94 variant.

Our work reveals an essential structural role of *N*-glycans in modulating the internal dynamics of the protein linked to the activation of functional motions. In particular, the carbohydrate moiety at N62 is shown to modulate the conformational states of the N-terminal domain (NTD) lid that controls the entry path to the ATP binding site, and to impact on the dynamics of the loop connecting the NTD and middle domain (MDomain), where client binding occurs. Deletion of different glycans results in different dynamic states. The simulations also show how two GRP94 inhibitors, PU-WS13 and PU-WS12, the first able and the latter unable to productively engage the disease-associated GRP94 variant, display a distinct cross-talk with the protein, leading to the selection of different dynamic ensembles of the protein. Overall, the data show that PTMs favor distinct states of the protein, which will ultimately be presented to different pools of interactors for the formation of assemblies that determine normal or disease cell phenotypes. The insights we provide here lay the foundation for the design of ligands that modulate GRP94 forms in a context-dependent way, which could be exploited in the development of drugs and chemical tools to untangle the intricate roles of post-translationally modified chaperones in disease. Furthermore, while focused on GRP94, the approaches we present and the models we develop here are entirely and immediately transferable to other systems, setting the stage for a generalized understanding of the interplay between protein covalent modification, the modulation of the conformational dynamics of specific structural elements (including active sites, disordered functional regions, client binding site etc.) and their impact on the large-scale motions associated to functions and interactions in cells.

## Results

### Ligand preference for GRP94 and disease variants: experimental investigations

GRP94 is a protein widely explored for the discovery of small molecule binders, and several GRP94 ligands were reported over the past few years. These include ligands based on the purine-scaffold (eg. N-ethylcarboximinoadenosine, cytosine-containing adenosine analogs, PU-WS13 and others) ^24–26^, resorcinol type compounds (eg. BnIm and KUNG94) and benzamide scaffold derivatives ^27–29^. These compounds, while tight binders of recombinant GRP94 protein, elicit non-overlapping biological profiles when tested in a variety of disease models. For example, some but not all are toxic to cancer cells characterized and driven by Receptor Tyrosine Kinase (RTK)-overexpression (eg. HER2- and EGFR-overexpressing breast cancer cells)^27,30,31^. In these cells, the N62 glycosylated GRP94 variant (referred to further as ^Glyc62^GRP94) is enriched at the plasma membrane, where it acts to reduce RTK internalization and to maintain the RTK in a state that is competent for constitutively enhanced downstream signaling^15^. Thus RTK-overexpressing cancer cells require ^Glyc62^GRP94 to promote survival, and its inhibition should be toxic in such context, consistent with previous findings for GRP94 knockdown in these cells ^15,24,32^.

The lack of biological activity observed for some GRP94 ligands but not others in RTK-overexpressing breast cancer cells has thus been puzzling. It was proposed, but never formally tested, that distinct ligands might favor and/or engage different context-specific GRP94 conformers and assemblies, each endowed with distinct biological function ^30^. To test this hypothesis, and thus provide molecular insights for atomistic studies through MD simulations, we selected a sub-library of closely-related GRP94 ligands of the purine-scaffold series (i.e. PU-series) (**Figure 1a**). We tested each ligand in a battery of assays designed to distinguish active ligands from those inactive on ^Glyc62^GRP94-related biological functions (see **Figure 1a-e** and **Supplementary Figure 1a-c**). We contrasted these ligands and their activity to BnIm, a ligand reported to productively engage GRP94 in cells, and modulate GRP94-but not ^Glyc62^GRP94-related functions^15,27,30,31^.

**Figure 1.**
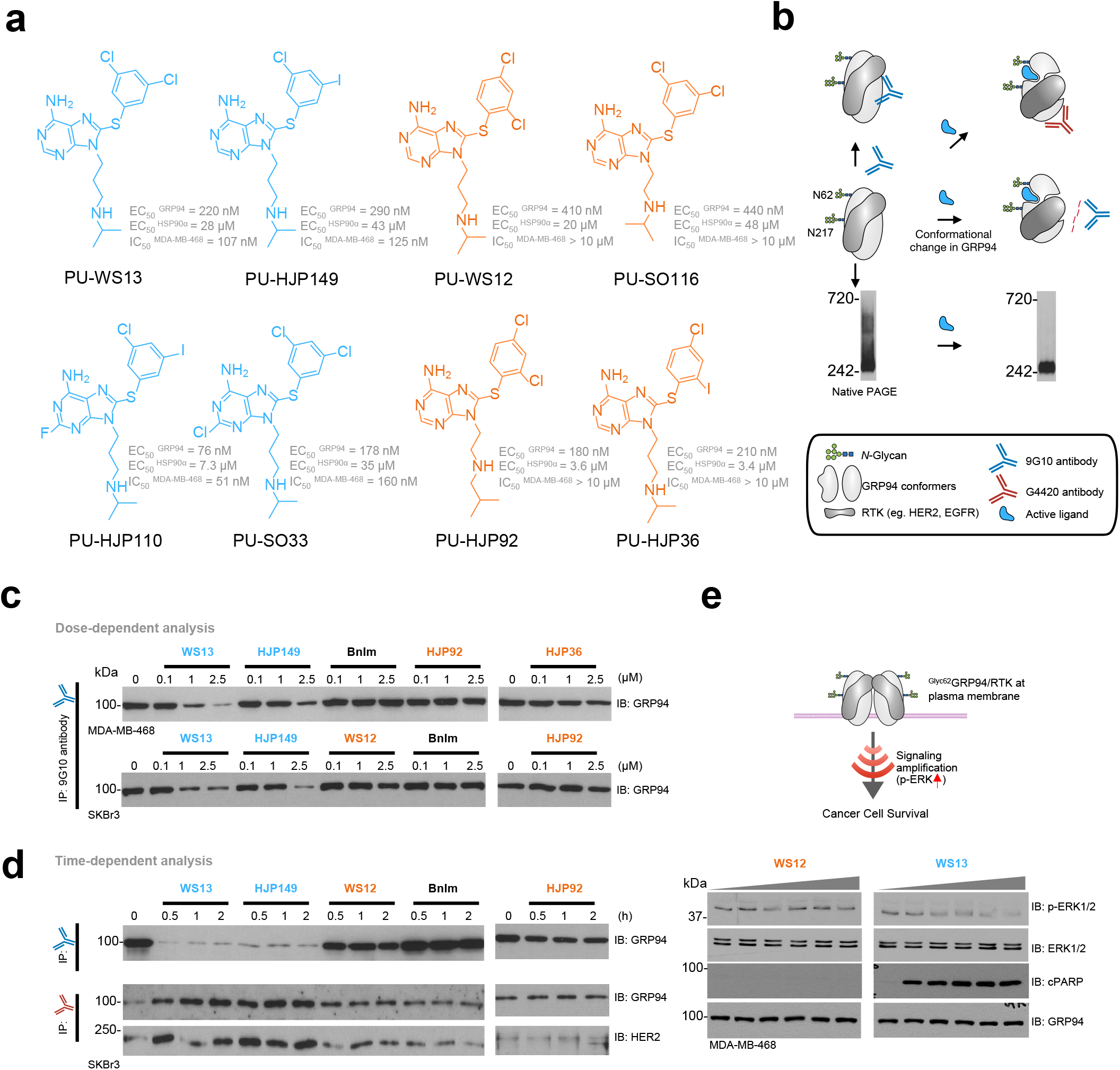
Biochemical and functional evaluation of GRP94 ligands for biological activity via ^Glyc62^GRP94. (a) Chemical structure of GRP94 small molecule ligands. EC_50_^GRP94^ and EC_50_^HSP90^, relative binding affinity constant to recombinant GRP94 and HSP90α, respectively. IC_50_^MDA-MB-468^, half maximal inhibitory concentration determined in the MDA-MB-468 EGFR-overexpressing and ^Glyc62^GRP94-expressing breast cancer cells (ATP quantitation assay). Values, mean of three independent experiments. (b) Schematic shows conformational changes in GRP94 expected from small molecule ligands that act on ^Glyc62^GRP94. The distinct GRP94 conformers are sensitive to detection by the indicated antibodies. (c,d) Immuno-capture as in (b) of SKBr3 HER2-overexpressing or MDA-MB-468 EGFR-overexpressing (both ^Glyc62^GRP94-expressing) breast cancer cells treated with the indicated ligands in a dose- (for 4 h) and time-dependent (at 2.5 μM) manner. Experiments were repeated thrice with similar results. (e) Western blot analysis for inhibition of ^Glyc62^GRP94 function in MDA-MB-468 cancer cells. Downstream signaling (p-ERK/ERK) and induction of apoptosis (cleaved PARP, cPARP) were analyzed for MDA-MB-468 cancer cells treated for 24 h with 0, 0.5, 1, 2.5, 5 and 10 μM of the indicated inhibitors. GRP94, loading control. Gels are representative of three independent results. See also Figure S1.

We found most ligands exhibit a comparable binding affinity for recombinant GRP94 under conditions of equilibrium binding (**Figure 1a**, see EC_50_ rGRP94, 120-450 nM for the PU-series and 1.2 μM for BnIm, as reported ^15,27,30,31^. Remarkably, even in the structurally closely related PU-type ligands, some but not all, were active in cancer cells dependent on ^Glyc62^GRP94 for survival, even at the highest tested concentration of 10 μM, which is 1- to 2-log higher than their recorded binding affinity constant (**Figure 1a**).

To provide mechanistic insights into the observed differences, we implemented a testing pipeline that probes biochemically and functionally productive engagement of ^Glyc62^GRP94. It is based on the observations that preferential and productive binding to ^Glyc62^GRP94 is dependent on a GRP94 conformation being stabilized by N62-glycosylation^15^. This conformation is sensitive to detection by GRP94 antibodies, such as 9G10, which recognizes the charged linker region (residues 290–350), and G4420, which recognizes amino acids 733–750 in the C-terminal domain of GRP94 (**Figure 1b**). We treated RTK-overexpressing breast cancer cells (eg. MDA-MB-468, EGFR overexpressor and SKBr3, HER2 overexpressor) with the ligands prior to immunocapture with the two GRP94 antibodies (**Figure 1c,d**). Active ligands, but none of those inactive, dramatically reduced the amount of GRP94 captured by the 9G10 antibody, indicating that active inhibitors change GRP94 to a conformation that is no longer recognized by the antibody. Active ligands also increased the amount of both GRP94 captured by the G4420 antibody and HER2 pulled down by the immunocapture of ligand-bound GRP94, indicating that active ligands promote or stabilize a conformation that favors HER2 association. Cell permeability or efflux pump mechanisms were not accountable for the activity difference (see activity in cell homogenates *vs* intact cells, **Supplementary Figure 1a**).

Native PAGE analyses confirmed the immunoprecipitation results (**Supplementary Figure 1b**). Under non-denaturing gel conditions, cells expressing ^Glyc62^GRP94 exhibit a number of distinct and indistinct high-molecular-weight GRP94 species above the main 242-kDa band (**Figure 1b**), a biochemical signature of ^Glyc62^GRP94 incorporation into epichaperomes, high molecular weight pathologic scaffolding structures formed of tightly bound chaperones, co-chaperones and other factors^15,33^. The signal observed in native PAGE reflects both GRP94 complexation and retained native conformation of the protein, which is recognized by the 9G10 antibody. We observed that compounds active in cells *via* ^Glyc62^GRP94, and not those GRP94-active but ^Glyc62^GRP94-inactive, induced a change in the epichaperome species detected on Native PAGE (**Supplementary Figure 1b-d**).

^Glyc62^GRP94 is required to reduce RTK internalization and maintain the RTK in a state that is competent for constitutively enhanced downstream signaling^15^. Compounds active in cells *via* ^Glyc62^GRP94, and not those GRP94-active but ^Glyc62^GRP94-inactive, inhibited RTK downstream signaling as measured by western blot analysis (see p-ERK, **Figure 1e**).

Overall, active ligands remodelled glycosylated GRP94’s conformation upon binding, as measured by antibody binding, HER2 capture, and native gel mobility, and inhibited ^Glyc62^GRP94 function in cells as measured by cell death induction and RTK function inhibition in ^Glyc62^GRP94-expressing RTK+ cancer cells (eg. SKBr3 and MDA-MB-468). Unlike active ligands, inactive ligands produced no or partial decrease in the conformation detected by 9G10 and G4420, no change in ^glyc62^GRP94-incorporating epichaperomes, and exhibited no cellular activity via ^Glyc62^GRP94. These findings are important in the context of our MD studies because they clearly show not every GRP94 ligand can optimally engage the ^Glyc62^GRP94 variant, but more importantly they indicate that we can study, and elucidate, the structure of ^Glyc62^GRP94 by investigating its dynamics and conformation in relationship to active and inactive ligands which we proceed to do below.

### MD simulations: design and implications for GRP94 functional investigations

The energy landscape model can help rationalize the fundamental physico-chemical mechanisms of how PTMs work. In this theoretical framework, biomolecules are not considered static objects but are instead viewed as dynamical entities that are continuously interconverting between a variety of structures with varying energies: distinct structures may be endowed with different functions depending on the conditions. Furthermore, by also describing the barriers between the different ensembles of states (inactive and active ones), a view on the kinetics of interconversion among structures and consequent functional regulation in the cell can begin to be understood. In this model, a PTM’s effect is both local and distal, as it modifies a residue and its immediate interaction networks, and importantly, it may propagate its effect throughout the structure, even at distant regions, eventually determining a protein population shift^2,3,8,9,34^. As a consequence, a PTM-modified protein may be devoid of enzymatic reactivity or show different mechanisms and affinities for ligand binding, while removal of the PTMs may shift the protein back to the catalytically competent structural ensemble, restoring enzymatic activity, ligand recognition mechanisms, and PPI interaction profiles^35–38^.

To understand how PTMs and inhibitor binding modulate the conformational dynamics of GRP94, we set out to analyze and compare fully solvated atomistic simulations of different models of GRP94. Specifically, we analyzed the protein with glycosylation at both N62 and N217 (simulations with double glycosylation, labeled 2Glyc), bound to the endogenous activator ATP (simulation labeled 2GlycATP) or to two different inhibitors, namely PU-WS13 (as a prototype active inhibitor, simulation labeled 2GlycPU-WS13) and PU-WS12 (ligand that binds but is functionally inactive on the tumor ^Glyc62^GRP94 variant, i.e. prototype of inactive inhibitor, simulation labeled 2GlycPU-WS12) (**Figure 2a,b**).

**Figure 2.**
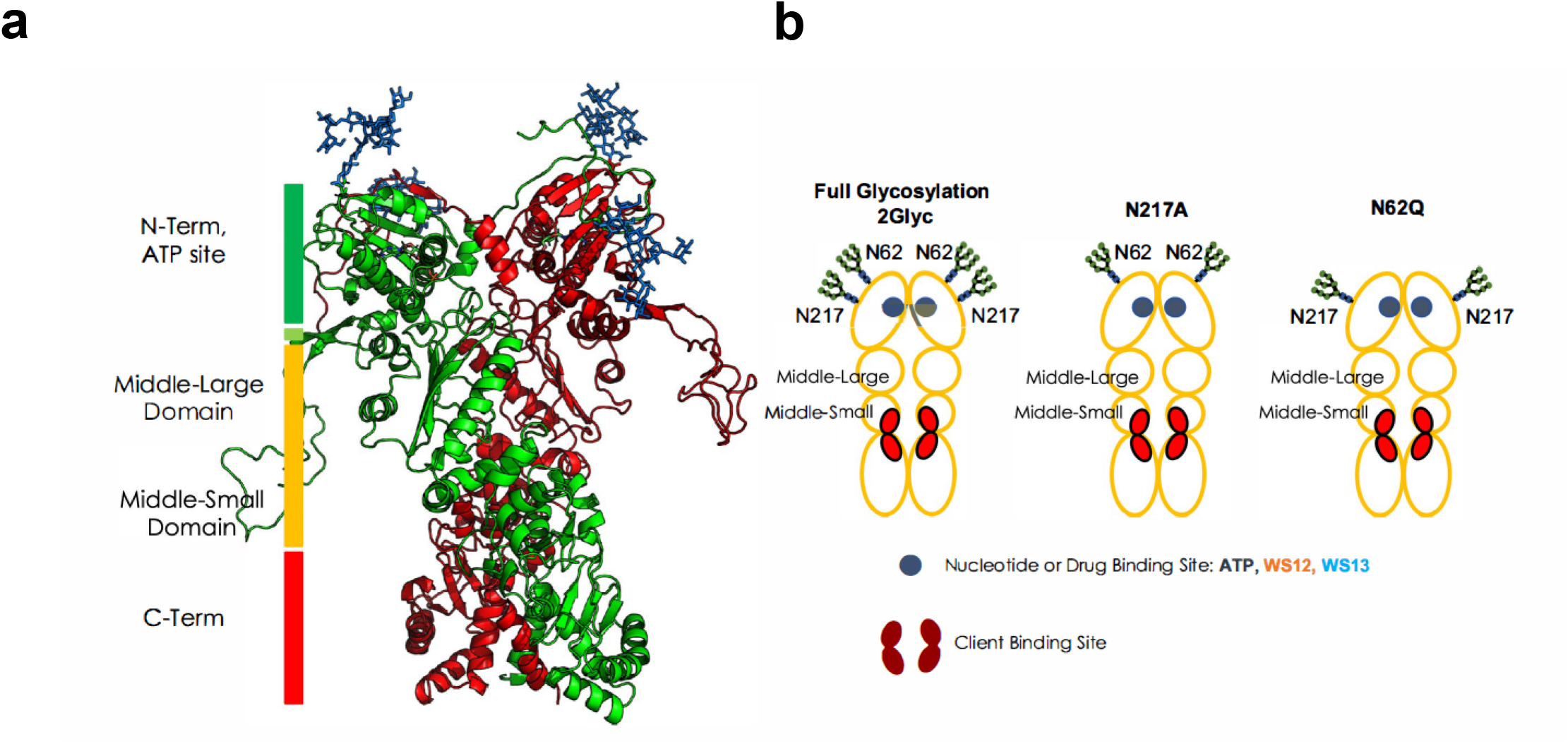
Structural and domain organization of GRP94. (a) The 3D structure of 2Glyc GRP94. The two protomers are colored in green and red respectively, while the glycosyl PTMs are shown in blue sticks. (b) Schematic of the domains, binding sites, glycosylation sites, and glycosylation states, as they are referred to throughout the paper. The red areas indicate the client binding site regions.

To understand the roles of the sugars in modulating functional dynamics, we generated two experimentally inspired mutants, namely N62Q (no glycosylation at position 62 while retaining the glycosylation at position 217; simulations labeled N62Q) and N217A (no glycosylation at position 217 while retaining glycosylation at position 62; simulations labeled N217A, mimicking the experimentally studied ^Glyc62^GRP94) (**Figure 2a,b**). Each mutant was simulated in the presence of ATP, PU-WS12, or PU-WS13 in the active site: the labeling for these states follows the logic described above for the 2Glyc case.

Overall, here we considered and cross-compared 9 systems. Each was simulated for 3 microseconds in total (3 independent replicates of 1 microsecond length each), for a total of 27 microseconds. Throughout the paper, we will use the fully glycosylated ATP state as the reference state. Moreover, as controls, we also simulated three completely unglycosylated systems (3 microseconds each), labeled Wild Type (WT): these models are intended to provide an additional control reporting on the dynamics of GRP94 in the absence of PTMs. In the three control simulations, GRP94 is in complex with ATP (labeled NoGlycATP), PU-WS13 (labeled NoGlycPU-WS13), PU-WS12 (labeled NoGlycPU-WS12). All simulations, their conditions, and the respective labels are reported in **Table 1.**

### Glycosylation modulates the dynamics of GRP94 and its specific functional substructures

To start quantifying the effect of glycosyl moieties on the dynamic states of the protein, we first visually analyzed the ensembles of equally interspersed conformations of each glycan sampled along each 3 μs metatrajectory for a total of (300) superimposed poses^17^. This representation returns a realistic view of the mechanisms by which sugars can modulate access to GRP94 interactors (**Figure 3a-c**).

**Figure 3.**
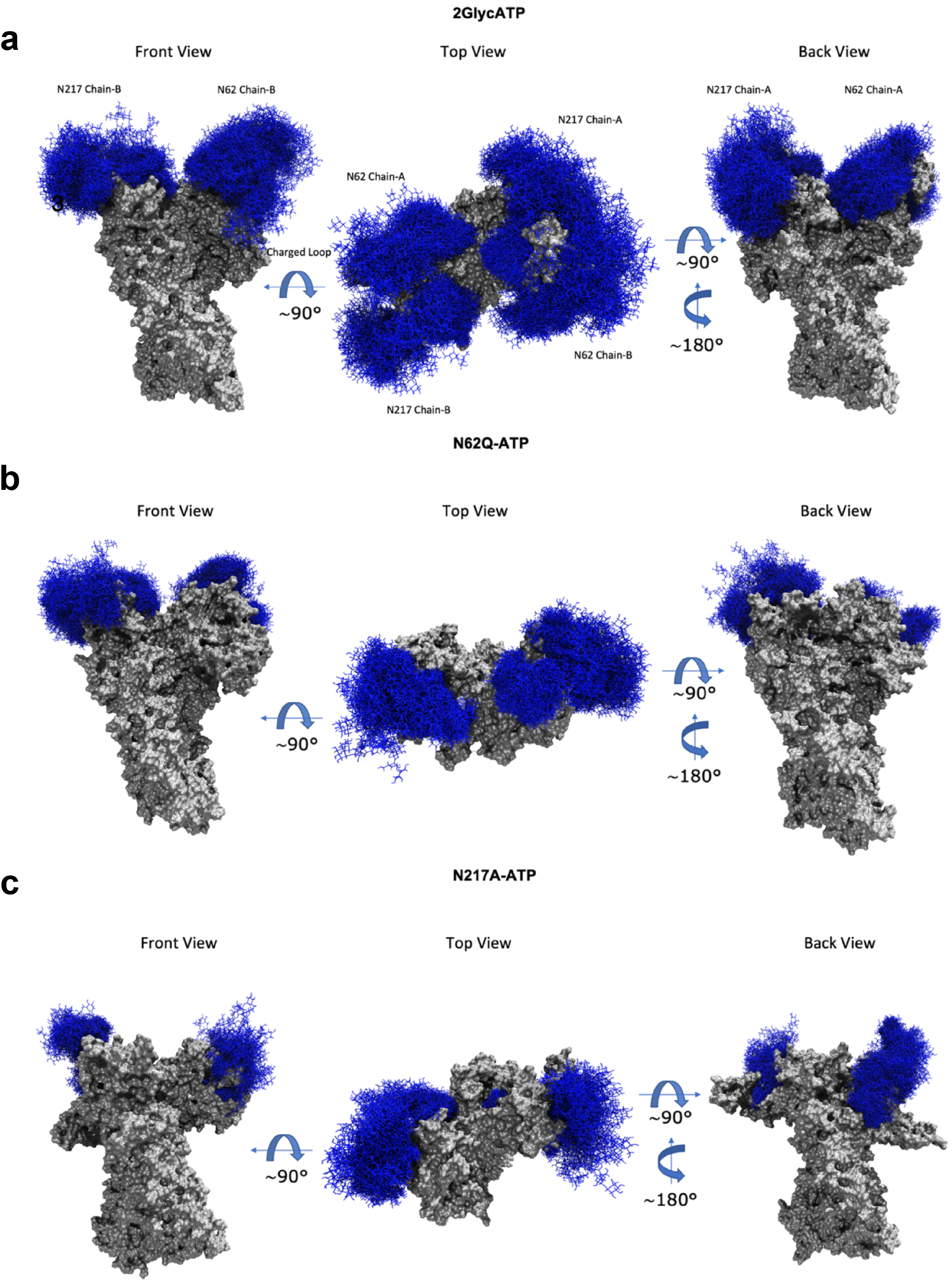
Glycan dynamics on the surfaces of GRP94. (a) Molecular representation of the 2GlycATP system. Glycans at several frames (namely, 300 frames, one every 30 ns from one replica) are represented with blue lines, while the protein is shown as a gray surface. (b) Same as (a), for the N62Q mutant in complex with ATP. (c) Same as (a), for the N217A mutant in complex with ATP.

In the ATP state of the fully glycosylated protein (2GlycATP), the sugars completely cover the NTD surface, with N-glycans at N62 extending out to form interactions with the long charged loop that connects the N-terminal domain (where ATP binds) to the M-domain where the client-binding site is located (**Figure 3a**) ^36–39^. Interestingly, removal of this glycan in the N62Q mutant removes the shielding from a large part of the NTD and releases the long N-M connecting loop (**Figure 3b**). In contrast, removal of N-glycans from N217 (N217A mutant, mimic of ^Glyc62^GRP94) with conservation of the extensive glycosylation on N62 further exposes the tendency for the latter to interact with the long loop (**Figure 3c**). The N-M loop plays an important role in regulating the conformational cycle of the chaperone upon ATP hydrolysis, connecting nucleotide processing in the NTD with structural reorganizations of the M-domain region. In our model the N-glycan chain at N62, the defining characteristic of ^Glyc62^GRP94, modulates the motions of the loop, blocking it into stable interactions (vide infra for the analysis of intramolecular contacts). The motions around the hinge between the N and M-domains can thus be perturbed with clear functional consequences, namely blockage of the foldase activity of the protein and promotion of its scaffolding activity. In this framework, our data also imply that the role of N-glycans at N217 is less relevant for functional regulation.

We next calculated the frequency with which a contact between the sugar chain and the protein is formed (see Supplementary Information Figure S2-S4). The full data are provided electronically (Data_Glyc_Contacts.zip, see Key Resource Table). In the fully glycosylated form, both glycosyl chains interact with the ATP-lid (see next paragraph too), from residue 168 to 181. Removal of glycosylation at N217 appears to increase the number of contacts of N62 glycosyl with the long charged loop: here, the interactions are established over the entire loop, from residue 280 to 330. Removal of glycosylation at N62 does not seem to affect the behavior of glycosylation at N217, with the glycosyl chain at this site acting as in the fully glycosylated form. It is important to underline here that the significant shielding by the sugars can furthermore disfavor binding of ATP to the NTD by rendering the nucleotide binding site less accessible, thus significantly decreasing the efficiency of GRP94 in the folding cycle.

The interactions that the carbohydrate chains at N62 establish with the N-M charged loop, a key factor in relaying the conformational signal encoded by ATP from the NTD to the M-Domain, can aptly perturb the structural dynamics of client binding site required for client-folding^40–43^. To investigate this point, we set out to characterize the deformations of the client binding site: we calculated the distribution of the Root Mean Square Deviation (RMSD) of client binding pockets extracted from the various trajectories from the structure of the client binding pocket determined by X-ray crystallography (PDB structure with code 5ULS^43^). This calculation, combined with the visualization of representative structures in **Figure 4a-d**, shows how different PTM conditions indeed differentially remodel the client binding pocket. In the 2GlycATP state, significant RMSD deviations can be observed for both protomers (peaks of the distributions at high RMSDs). From the structural point of view, a closure of the space between the two protomers at the level of the client-binding site can be noticed. In contrast, in the NoGlycATP state, the intraprotomer space in the lumen at the client binding site is substantially larger and the RMSD distributions are centered on lower values. In the fully glycosylated form, the reduction of the intraprotomer space hinders recognition, binding, and the threading of the client through the lumen. All three processes would be disfavored compared to the NoGlycATP state, where the lower steric hindrance of the binding pocket can support adaptation to and conformational maturation of the client. Interestingly, the N217A mutant (^Glyc62^GRP94), while displaying a slight opening of the pocket, never reaches the states observed for unglycosylated WT.

**Figure 4.**
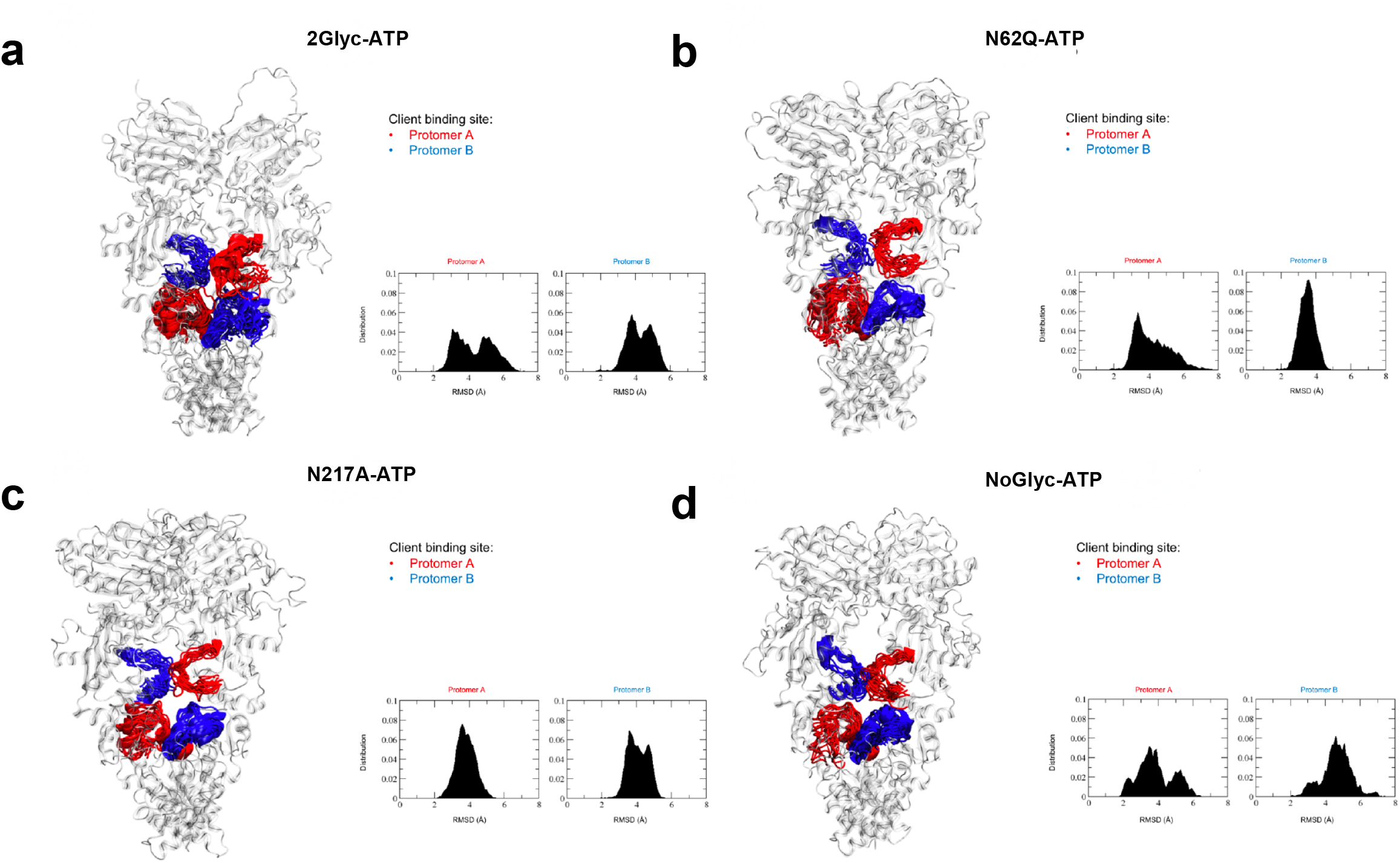
Conformational dynamics of GRP94 Client-Binding Site. (a) The 2GlycATP system. (b) Same as (a), for the N62Q mutant in complex with ATP. (c) Same as (a), for the N217A mutant in complex with ATP. (d) Same as (a), for the completely unglycosylated NoGlyc state in complex with ATP. The client binding site is depicted as red (Protomer A) or blue (protomer B) ribbons, whereas the rest of the protein is depicted as transparent cartoons. The insets show the distributions of RMSD values from the conformation presented in crystal structure PDB 5ULS.pdb.

Overall, these results lay the foundation for a mechanistic model in which glycosyl modifications play an active role in modifying the conformational landscape of GRP94, switching its functions from a foldase protein to a protein-assembly platform: specifically, glycosyls perturb the communication between the NTD and the M-Domain, shield the ATP-binding site in the NTD domain lowering the efficiency in binding and processing ATP, and favor the closure of the lumen through which the client needs to thread to be properly remodeled.

### Glycans and ligands differentially modulate the internal dynamics of GRP94 variants

To gain further insight into the impact of glycosylation on inducing dynamic changes and modulating ligand effects, we next computed various simulated complexes (see Materials and Methods). In the energy landscape framework, the effects of glycosylation, even if localized at a specific protein site, can diffuse throughout the structure and be distinctly sensed by groups of distal residues in functionally relevant regions of the protein ^44–46^. The DF parameter reflects the degree of internal coordination of residue-pairs in a structure and can consequently report on the diffusion of dynamic effects ^44,45^. Differences between the DF profiles of the reference state (in this case the fully glycosylated ATP state, 2GlycATP) and differentially glycosylated as well as inhibitor-bound states can identify regions that are responsive to the modifications. Here, we aim to investigate how changes on short time-scales in the structural dynamics of GRP94, captured by DFs, can cooperate in modulating the motions of regions that determine biological functions. Our working hypothesis is that distinct collective GRP94 fluctuation patterns that emerge as responsive to a particular ligand/glycosylation state may underpin such key functional motions. The modulation is achieved through the local disruption and reassembly of interactions, which will eventually result in specific structural deformations and modifications of the protein energy landscape on longer time scales ^2,7,47–50^.

The representative pairwise mean-square distance fluctuations DF matrix for the reference state 2GlycATP is reported in a color-coded form in **Figure 5a.** For the sake of brevity and simplicity, the fully detailed matrices are reported in the Supplementary Information in the Figures S5-S6. In **Figure 5,** we report a simplified view of the GRP94 domains whose internal dynamics is mostly perturbed compared to the 2GlycATP state according to the DF metric, as a consequence of glycosylation modification or ligand binding. This representation reflects the difference DF matrices reported in Supplementary Information Figures S5-S6.

**Figure 5.**
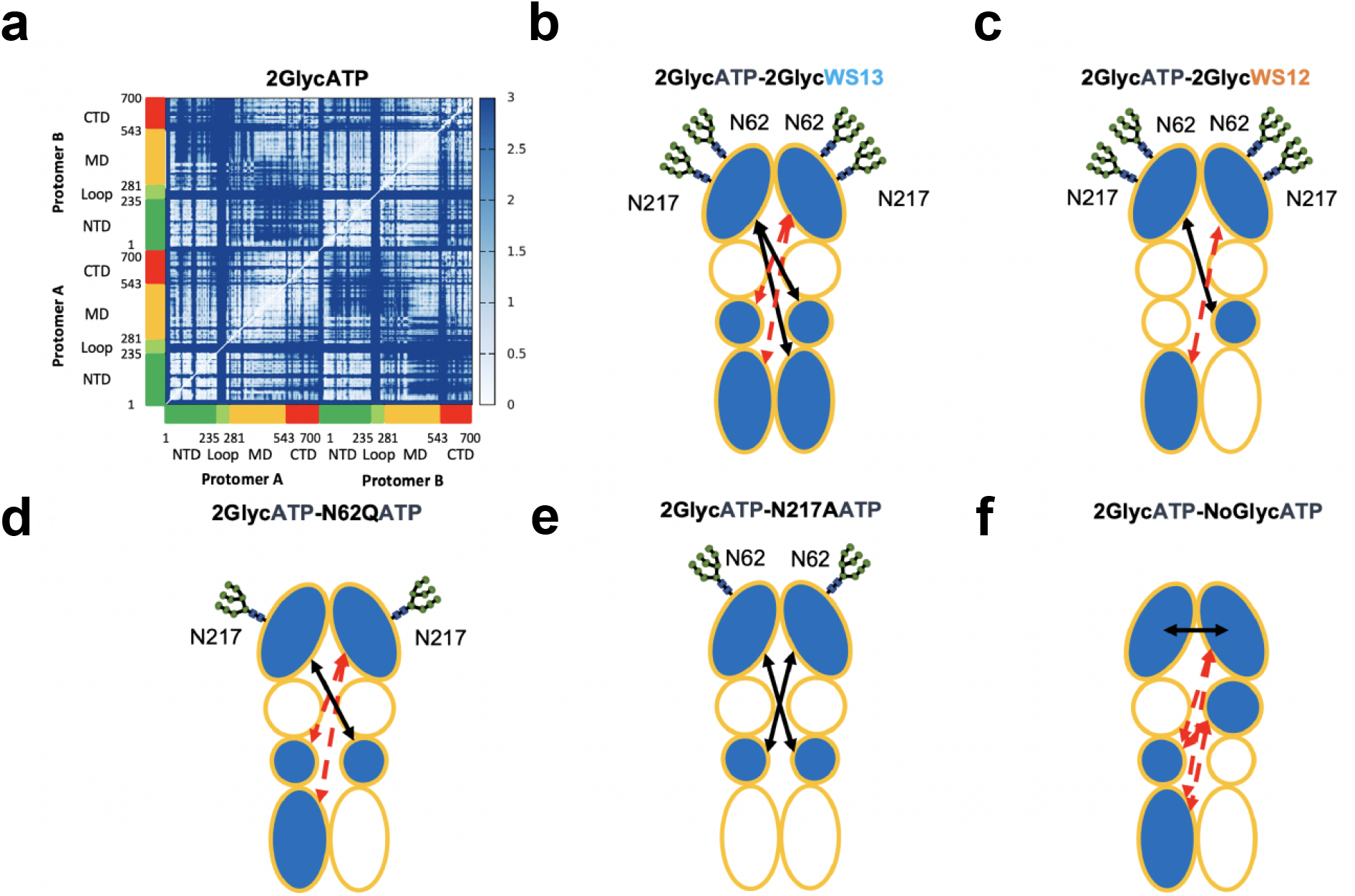
Residue-Pair Distance Fluctuations and Domain Cross-Talk for GRP94 in Different Glycosylation and Ligand Conditions. The internal dynamics of GRP94 characterized from the various trajectories. (a) The original Distance Fluctuation (DF) matrix is reported for the fully-glycosylated ATP-bound state. The axes report the residue numbering and domain organization, as depicted also in Figure 2. In this view, lighter pixels correspond to highly coordinated residue pairs, while darker ones report on low coordination pairs. The single original matrices for all 9 conditions, together with the difference matrices obtained by subtracting the DF matrix of a certain state from the reference one pertaining to 2GlycATP are reported in Supplementary Information. (b-f) Domain based representations of the modulation of internal flexibility as a function of ligand-state and/or glycosylation state. This representation is a simplified graphical translation of the difference matrices reported in the Supplementary Information (see Figures S5,S6). A certain domain is colored in light-blue if its coordination with other domains changes as a function of the condition. The variation in internal dynamics is evaluated as a difference for the DF of the system under exam from that of the fully glycosylated ATP-bound state, 2GlycATP. A black arrow indicates increased coordination (rigidity) between two domains with respect to the fully glycosylated ATP-bound state, 2GlycATP. A red, broken arrow indicates decreased coordination (flexibility) between two domains with respect to the fully glycosylated ATP-bound state, 2GlycATP. The various ligand states are reported in each subfigure.

From a general point of view, all matrices reported in SI exhibit a common block character, which reflects the alternation of regions with small and large fluctuations of inter-residue distances consistent with the tripartite domain organization of GRP94 protomers, the way different domains are coordinated with each other, and their relative flexibilities. As discussed hereafter, differences emerge in analyzing the finer details of the matrices which point to glycosylation-dependent modulations of the internal dynamics in different complexes. Relevant differences are also seen in the comparison between the active and inactive ligands.

We find that the fully glycosylated ATP (2GlycATP) state is characterized by extensive regions of coordination between the NTD, M-domain, and CTD within each protomer, between the NTD of one protomer and the NTD and M-domain of the other, and between M-Domains (**Figure 5a**)(See Supplementary Information Figure S5). In the PU-WS13-fully glycosylated GRP94 complex (2GlycPU-WS13), an increase in the fluctuations of pairwise distances can be noticed (large red areas in the difference matrix)(See Supplementary Information Figure S5). This entails in particular the relative fluctuations between the NTD of protomer 2 (chain B) and the Middle-Small/CTD substructure (which hosts the client binding site) of protomer 1 (chain A) compared to the ATP one (**Figure 5b**). The reported effects on the internal dynamics of the fully glycosylated version of the protein can be correlated with the observed efficacy of PU-WS13 in targeting this form in tumor models expressing it: the ligand in fact significantly perturbs the internal dynamics of the protein. In turn, the population of different dynamic states induced by the inhibitor can remodel the conformational fitness of GRP94 to interact with the set of proteins (such as RTKs) characterized in tumor cells where the connectivity of the fully glycosylated chaperone is a determinant of pathogenic dysfunction^15^. Importantly, these effects are significantly less pronounced as PU-WS12 is bound (Simulations 2GlycPU-WS12, **Figure 5c**)(See Supplementary Information Figure S5).

Removal of the N-glycan on N62 via N62Q mutation is seen to cause a decrease in the degree of coordination between the NTD of protomer 2 and the M-small/CTD region of protomer 1, showing DF patterns for N62Q-ATP state that are similar to the ones observed for the inhibitor-bound fully glycosylated complex (**Figure 5d**)(See Supplementary Information Figure S6a). In general, the presence of the synthetic ligands (in particular of the active PU-WS13) is characterized by a more marked effect on the internal dynamics and coordination patterns. Removal of the glycan on N217 (N217A mutant) in the presence of ATP, appears to have a much less pronounced effect than N62Q when compared with the 2GlycATP situation(See Supplementary Information Figure S6a). Indeed, the effect of maintaining N62, in terms of internal coordination, keeps the protein in a state qualitatively similar to that of the reference state (**Figure 5e**)(See Supplementary Information Figure S6a).

Importantly, these data are qualitatively consistent with experiments showing that N62Q mutant is inactive in tumor models, mimicking the effects of full GRP94 Knock Out or PU-WS13 inhibition. Similar patterns are observed as the two synthetic ligands are bound to N62Q or N217A (**Figure 6a-f**)(See Supplementary Information Figure S6b,c): the small molecules in both mutants modify mainly the internal coordination of the NTDs. Interestingly, PU-WS13 perturbs the intraprotomer dynamics (i.e. the coordination between the NTD with the M-Domain and the CTD) more than PU-WS12, compared to ATP.

**Figure 6.**
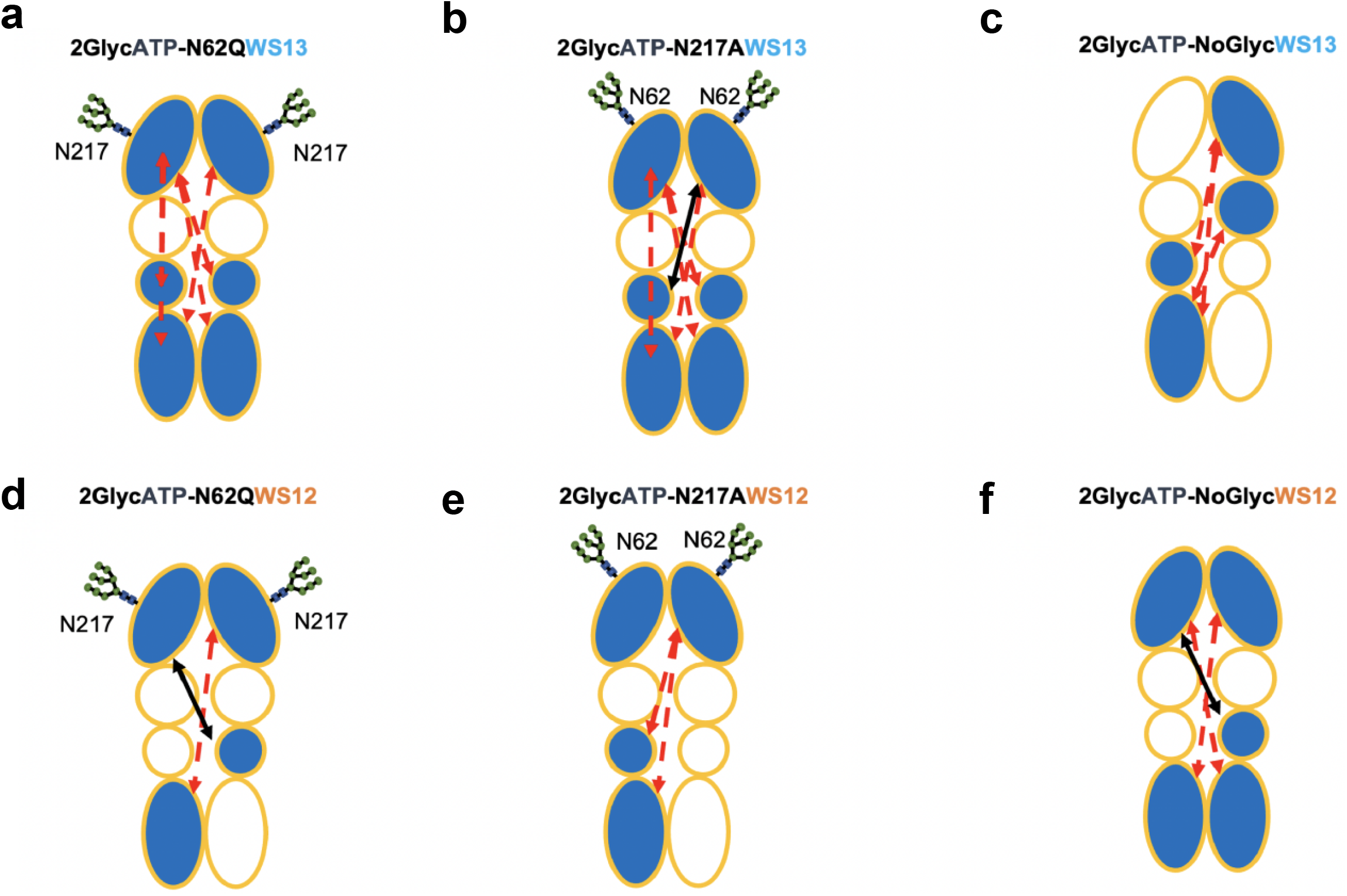
Residue-Pair Distance Fluctuations and Domain Cross-Talk for GRP94 in Different Glycosylation and Ligand Conditions. Same as Figure 5(b-f), for systems with different degrees of glycosylation. See also Supplementary Information, Figures S5,S6.

Interestingly, the fully unglycosylated WT system in the ATP-state (NoGlycATP) shows the high coordination between the two NTDs required to form the catalytically competent state ^45,51^. Significant flexibility is observed at the level of the Middle/CTD substructures (**Figure 5f**)(See Supplementary Information Figure S6a). These distinct dynamic traits of the NoGlycATP state can be reconnected to the foldase role of this GRP94 form. First, the closed-catalytic state is required for the proper organization of the ATPase enzymatic machinery; moreover, in a sequential model of ATP hydrolysis, the high coordination between the two NTDs favors sensing of the first ATP hydrolysis by the domain containing the second ATP molecule; finally, the flexibility of the client binding region supports the conformational changes that are required to adapt to the structures and carry out the remodeling of the client proteins^51^.

Overall, these data show that PTMs remodel the connectivity and coordination between distal substructures (sectors) of the protein that are required for proper functioning in normal conditions. Interestingly, PU-WS13 has a strong effect on the internal dynamics of protein variants with glycosylation at N62, a trait that can help explain the activity of this drug on highly glycosylated GRP94. In contrast, the impact of PU-WS12 is much less pronounced, which correlates with the apparent inactivity of this ligand.

### Ligands and PTMs modulate the conformations of the active site lid and of the nucleotide binding site

As a final step, we characterized the fine details of the interactions of glycosyls and ligands on the conformational profiles of the active site. The active site, and the region around it, is known to undergo large conformational variations, which entail open to close state transitions and modifications of the folded state of Helix 4 (H4), the main structural motif of the active site lid (**Figure 7**) reports three different conformations for the lid via a specific color code that labels distinct structures^43,52,53^.

**Figure 7.**
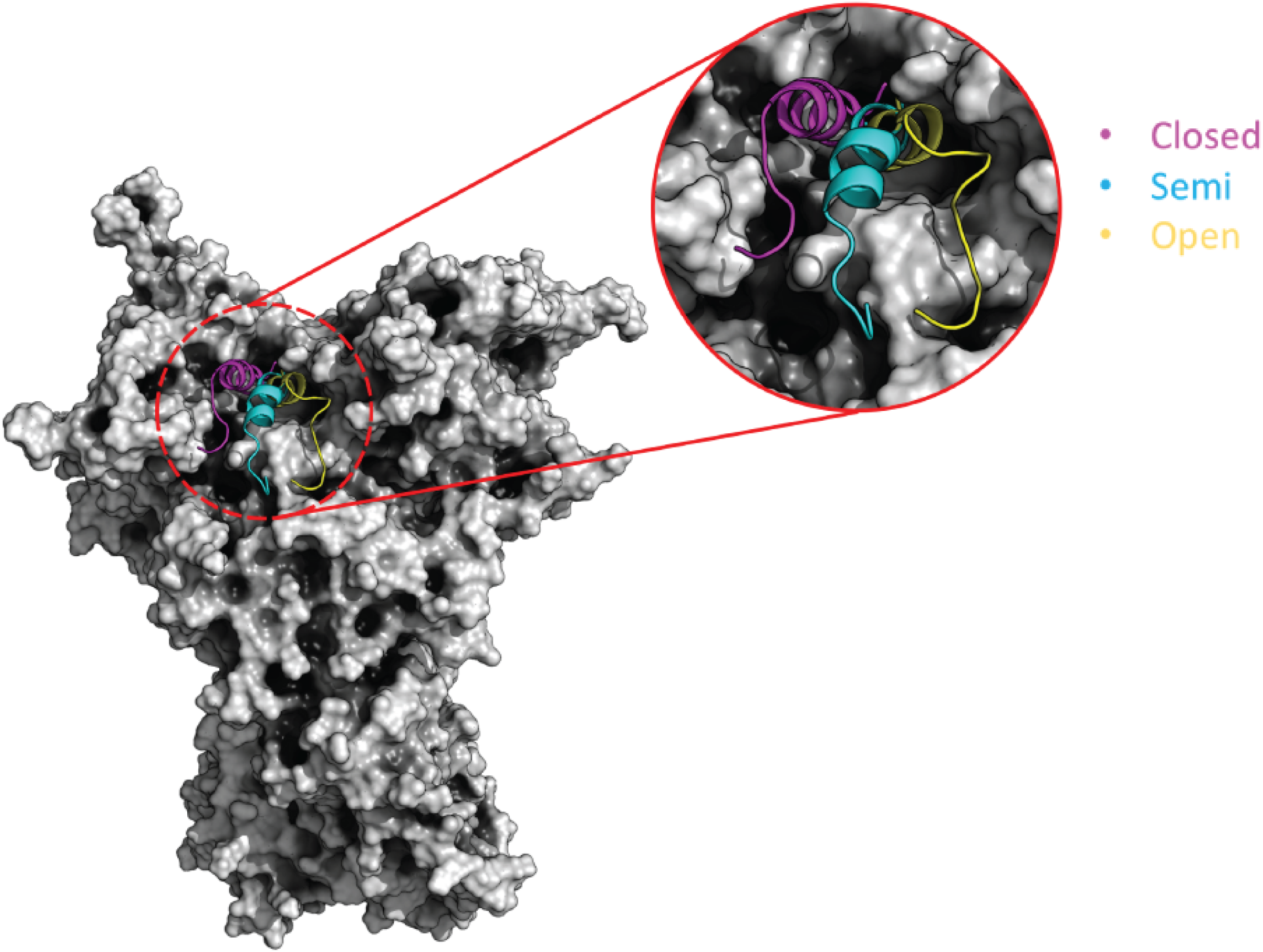
Simplified representation of structural ensembles of the GRP94 ATP-lid. The cartoon shows the whole GRP94 represented as a surface. The zoom on the lid structure shows the three main conformational families of the lid covering the active site of GRP94 in the closed (magenta), semi-open (cyan), open (yellow).

Simulations start with the lid in the closed state over the binding site. In the fully glycosylated state, in the presence of ATP (2GlycATP), the lid passes through closed and open states (**Figure 8a**). The figure shows in red the interspersed conformations of the lid sampled along the 3 μs metatrajectory for a total of (300) superimposed poses and in blue sticks the conformations of the sugars interacting with it. The inset, showing the distribution of Root Mean Square Deviations (RMSD) calculated with respect to the initial closed conformation of the lid for each protomer, indicates how the glycan on N62 stabilizes the open lid, in particular in protomer A, through interaction with helix H4 (**Figure 8a**). As the interaction is lost (N62Q mutant bound to ATP, **Figure 8b),**the distribution of lid conformations for protomer A shifts to the preferentially closed state. Here, the carbohydrate moiety bound to N217 folds on H4 stabilizing the closed conformation. Deletion of N217 (N217A mutant bound to ATP, **Figure 8c**) facilitates the population of an intermediate state for both protomers, with the ATP-lid of protomer B being still preferentially closed.

**Figure 8.**
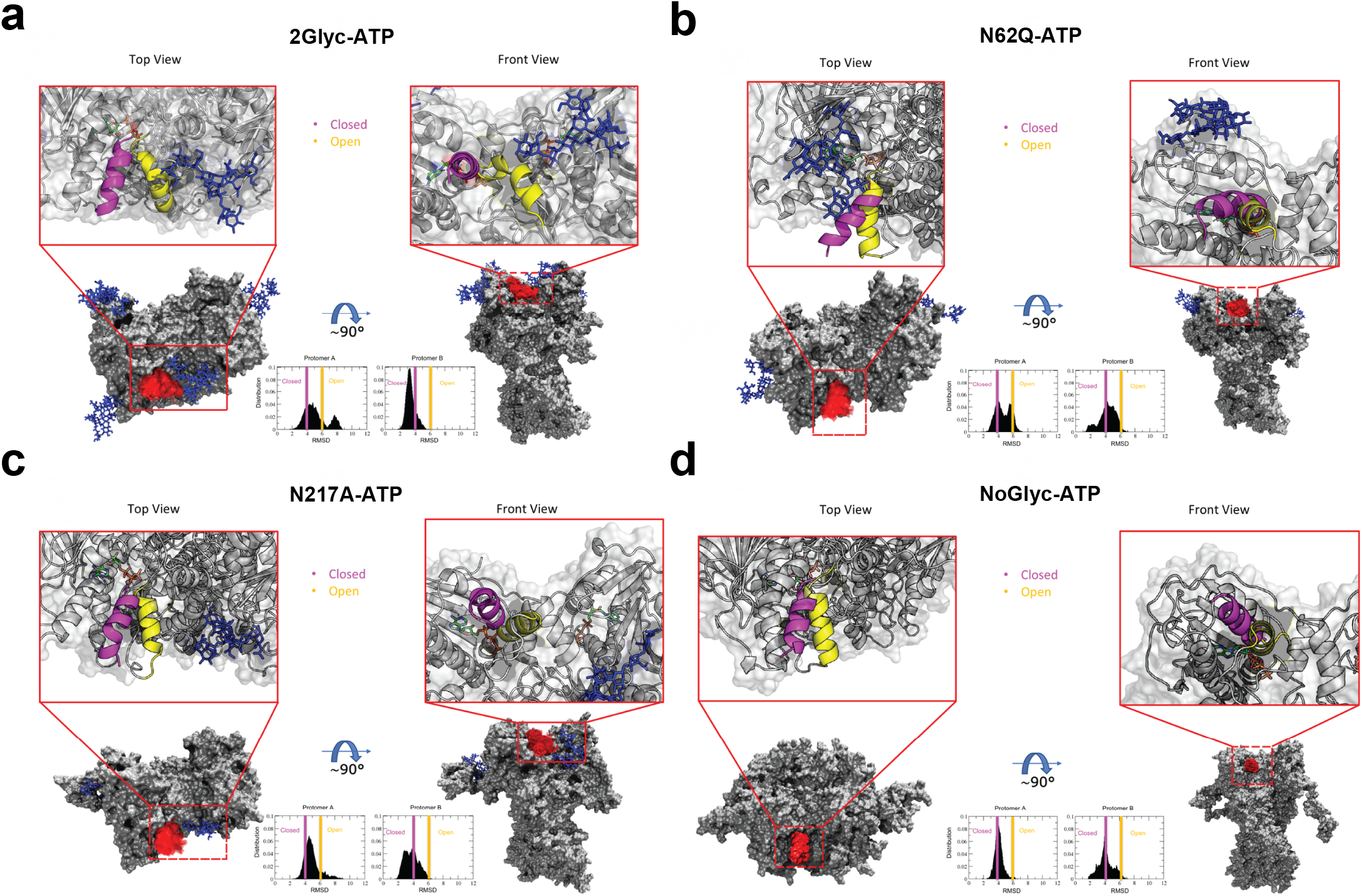
Conformational dynamics of the ATP-lid covering the active site of GRP94 and its interactions with sugar chains in the open or closed forms. (a) 2GlycATP. (b) N62Q mutant in complex with ATP. (c) N217A mutant in complex with ATP. (d) NoGlyc system in complex with ATP. Two orientations are depicted. ATP-lid and glycans at several frames (300 frames) are represented with red and blue lines, respectively. The insets show the distributions of RMSD values from the conformation presented in crystal structure PDB 5ULS.pdb. The blue and yellow vertical lines show the RMSD threshold for the closed and open states, respectively.

Overall, in the presence of the endogenous ligand ATP, the dynamics of the lid are regulated mainly by the glycan on N62, which stabilizes the open structure. As the closed lid-state is induced in the presence of ATP, the glycan on N217 may play an accessory role in stabilizing this conformation. Indeed, N217A mutation does not completely abolish the population of the open state. In contrast, as N62 is mutated to Q, the percentage of open lid conformations, in particular in protomer A, is significantly decreased.

To confirm the role of the glycans, we characterized the structural behavior of this ATP-lid in the unglycosylated form (**Figure 8d**). The difference in the selection of conformational ensembles compared to the cases reported above is striking, showing a clear preference for the closed state, and confirming the active role of PTMs in the modulation of the dynamics of functionally important substructures.

To shed light on the impact of a GRP94 small molecule ligand with high affinity for the fully glycosylated form, we analyzed the conformational dynamics of the lid in simulations with PU-WS13 bound to the active site (**Figure 9a)**. In these simulations, a clear partial unfolding of Helix H4 in the lid immediately emerges. In parallel, the long disordered loop in the lid is remodeled into a helical conformation, a variation that is not observed in the simulation with ATP. In these three replicates, the opening and closing of the lid is observed more frequently than in the simulations with ATP. The N62 glycan stabilizes the open lid structure, as observed previously in the ATP simulations. The Pre-N-terminal (PN) strap also participates in this interaction. In the presence of PU-WS13, the N217-glycan can act as a stabilizer of the closed form of the H4 of the lid.

**Figure 9.**
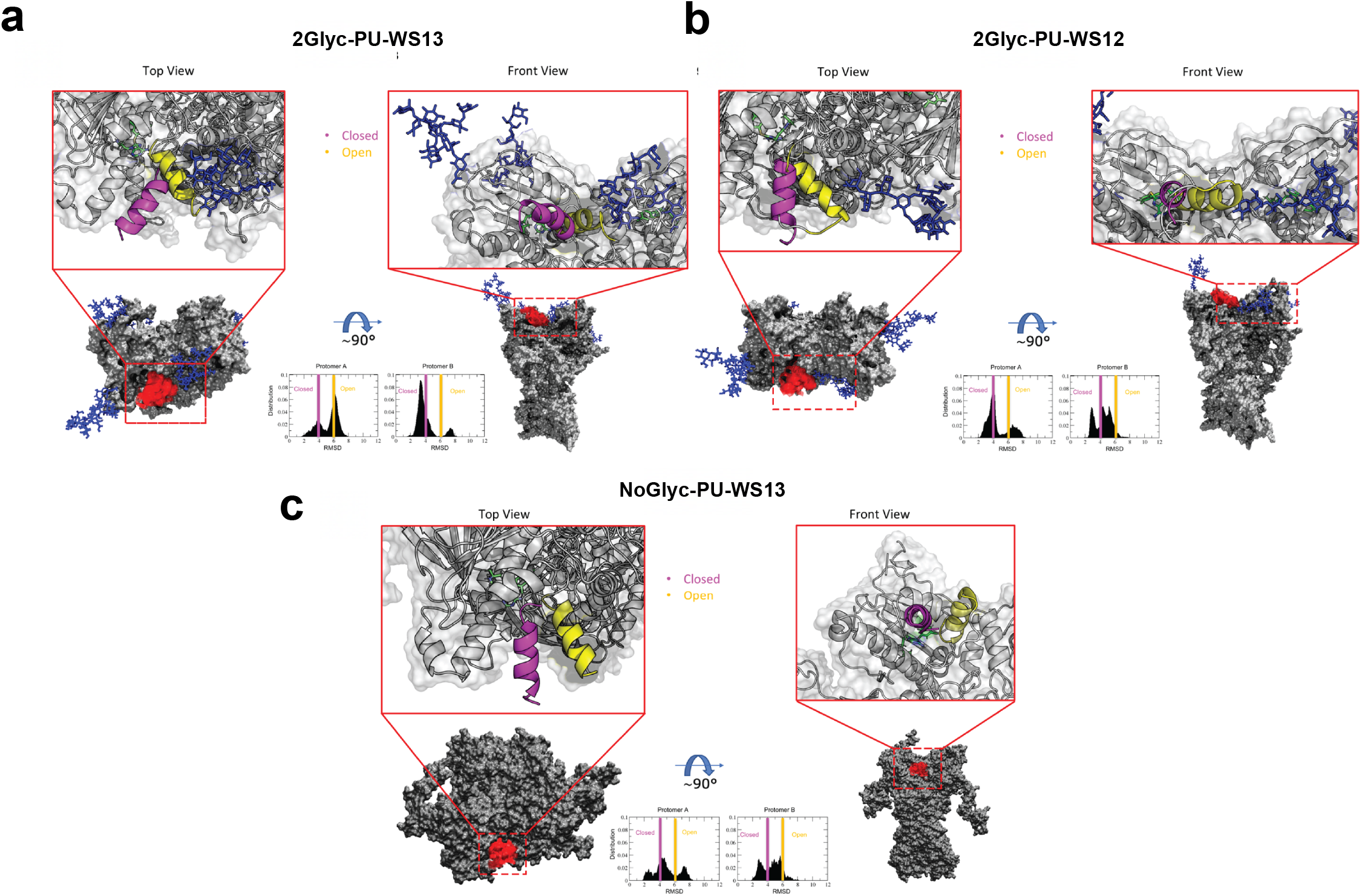
The effect of ligands on the conformational dynamics of the ATP-lid covering the active site of GRP94 and its interactions with sugar chains in the open or closed forms. Same as Figure 8. (a) 2Glyc system in complex with PU-WS13. (b) 2Glyc system in complex with PU-WS12. (c) NoGlyc system in complex with PU-WS13.

In summary, PU-WS13 reconfigures the structure of the lid, making it more dynamic and prone to spontaneously open. The effect of the glycans appears to be less pronounced than in the ATP-bound case, although the interactions that they establish are substantially the same as observed above.

In N62Q (Simulation N62Q-PU-WS13, See Supplementary Information Figure S7a), PU-WS13 induces a significant remodeling of the lid, which shows an increased tendency to populate an open conformation. No interactions with the lid for the N217 glycan are observed, a factor that would further favor an open configuration of the binding site. On this basis, it may be hypothesized that the residence time of PU-WS13 at the binding site would be significantly lower than that observed for the fully glycosylated form: a consistently open site can aptly facilitate the escape of the ligand from the protein. This model is consistent with the observed insensitivity to PU-WS13 of N62Q mutant cells ^15^. Importantly, the dynamics of the lid in the complex with PU-WS13 and the N217A mutant of GRP94 are qualitatively similar to the fully glycosylated GRP94/PU-WS13 complex (See Supplementary Information Figure S7b). This observation, in turn, is consistent with the fact that there is no effect on the binding of this ligand in the N217A mutant.

Finally, when PU-WS12 is bound, the lid appears to be partially unfolded as in the simulations with PU-WS13. The lid undergoes opening/closing transitions with and without interactions with the glycans. Also in this case, the presence of the ligand confers more mobility to the lid than is observed with ATP (**Figure 9b**). The glycan on N62 stabilizes the open form and the closed one can in some cases be stabilized by the glycan on N217 (**Figure 9b**)(See Supplementary Information Figure S8a,b).

In the NoGlyc form both ligands stabilize the open state of the lid, with PU-WS13 strongly favoring the open form: the impact of PU-WS13 in preventing the protein to stabilize the closed state required for catalysis and to enter the chaperone cycle is one of the factors that determine the efficacy of this molecule in tumor treatment (**Figure 9c**) (for NoGlyc-PU-WS12 See Supplementary Information Figure S8c)

Focusing on the ligand-binding site, we next calculated the frequency with which a contact is formed between the ligand and the surrounding residues of the protein: We report the results in a pictorial manner in the Supplementary Information Figure S9. In particular, we subdivided the residues around the ligands based on their frequency of contact with it. In yellow we represent residues that make contact with the ligand 20% to 50% of the time, orange 50% to 70%, red 70% to 100%. These data are also reported in a compact Table representation supplemented electronically (Data_Ligand_Contacts.zip, see Key Resource Table). Differences were observed as a function of the ligands. In the fully glycosylated protein, ATP exhibits the largest number of highly persistent contacts compared to PU-WS12/13. The low-activity ligand PU-WS12, in particular, appears to populate different ensembles of conformations in the complex, establishing in general mostly transient contacts with the binding site. These data also support a model whereby PU-WS13 features a longer residence time on the glycosylated target, determining a favorable kinetic profile.

These observations are further corroborated by analyzing the distributions of RMSD values of the residues making up the active site compared to their respective positions in the ATP-bound crystal structure of GRP94 (see Supplementary Information Figure S10). It is immediately evident that, in the fully glycosylated protein, the active site in the presence of ATP preferentially visits one ensemble of conformations, for both protomers. The same is noticeable for PU-WS13, which clearly locks the active site in a restricted portion of configurational space. PU-WS12, in contrast, explores a larger ensemble of configurations (larger distributions, with more than one peak). Importantly, removal of the glycosyl moieties in the WT system, determines markedly different profiles: a configuration similar to the one observed in the crystallographic closed structure is predominantly populated only for the ATP state in protomer A, while the structure of the binding site in protomer B appears to be endowed with larger conformational freedom. The sequential and deterministic model for ATP hydrolysis (proposed by Agard and coworkers)^51,54,55^ provides a rational framework for this observation. ATP hydrolysis occurs with different rates in the WT two protomers: while protomer A is ready to favor the reaction of the nucleotide, the other protomer needs to be flexible and explore different possible rearrangements for the second hydrolysis. Both PU-WS13 and PU-WS12 determine a variation in the RMSD distributions: interestingly, PU-WS13 is the ligand that forces the WT system to differentiate more substantially from the ATP-bound state.

In summary, the comparative analysis of the lid and active site dynamics defines a model in which the specific propensity of the GRP94 ATP-lid to populate open conformations is further amplified by the interactions established in particular by the glycosyl moiety at N62. Favoring the open state of the lid contrasts the population of the ATPase competent state. In this framework, the results are consistent with the experimental characterization of N62-glycosylation as a key factor for the switch of GRP94 from a folding ER chaperone to a templating protein. Interestingly, the active ligand PU-WS13 further favors lid opening and unfolding of H4 in the N62 glycosylated versions of the protein. At the same time PU-WS13 populates one dominant configuration in the active site, establishing stable interactions with it. Qualitatively speaking, this can be expected to determine a longer persistence time of this ligand compared to PU-WS12, providing further mechanistic support to the observed activity of PU-WS13 in cells characterized by aberrant GRP94 glycosylation profiles.

### Binding of different scaffolds: GRP94 conformational preferences underlie the selection of active ligands

Several GRP94 ligands with different structures and scaffolds were reported over the past few years. These include for instance resorcinol type compounds such as BnIm, which are GRP94 active but ^Glyc62^GRP94 inactive (see **Figure 1** and **Supplementary Figure 1**) ^28,29^. Here, we tested whether these types of compounds could preferentially bind to protein conformations representative of different ensembles. This test is also intended to assess if the conformations obtained from the simulations are actionable in the selection of ligands for distinct protein forms. To this end we docked BnIm ^28^ into the active site of the central structures of the most populated conformational clusters obtained from the trajectories of the fully glycosylated and the WT forms bound to ATP, respectively 2GlycATP and NoGlycATP (see Supplementary Information Figure S11)(Full data are provided electronically as Data_Docking_Bnlm.xlsx, see Key Resource Table). Interestingly, while in the case of NoGlycATP the best docking pose is observed for Bnlm binding to the most visited conformation, in the fully glycosylated 2GlycATP form, the best docking poses are observed for the states that are visited less frequently. This result is consistent with the fact that BnIm ^28^ was experimentally shown to be unable to inhibit glycosylated GRP94, as evidenced biochemically (see the effect on GRP94 conformation, **Figure 1c,d and Supplementary Figure 1c,d**) and functionally (no cytotoxicity and no effect on HER2 signaling in SKBr3 cells at the highest tested concentration of 20 μM, 1 −log higher than its binding affinity for GRP94)^30^.

Summarizing, our data suggest a model whereby PTMs and ligands cooperate in defining different GRP94 dynamic states. In particular, the observed modulation of the active site provides a structural framework to rationalize the distinct effects of the ligands on the various PTM-modified forms of GRP94, and, as a consequence, their impact on the biology of cells where these forms are highly expressed.

## Discussion

Our study unveils the mechanisms through which *N*-glycans at different protein sites modulate the conformational dynamics of GRP94, affecting both its nucleotide processing activity and interaction modes with clients, at atomistic resolution. Specifically, we build a structural and dynamic model in which the degree of glycosylation and the specific positioning of the glycosyl moieties are correlated with the observed switch of function for the PTM-modified GRP94 from a folding to a scaffolding protein. In our model, stress-induced glycosylation, in particular at position N62, favors dynamic states of the chaperone with peculiar traits of internal coordination, distinct structures of the client-binding site, and specific responses to nucleotide and inhibitor binding.

Our data support an active role for glycosyl modifications: consistent with experimental observations^15^, in the presence of glycosylation at N62 (fully glycosylated protein or N217A mutant) the combined perturbation of the communication between the NTD and the M-Domain, the expected lower efficiency in binding and processing ATP due to sugar-shielding, and the closure of the lumen can block remodeling of the client^15^. In this context, glycosylation affects the conformational modifications of the chaperone and its ability to relay the signal encoded by ATP to the client binding site^20,56^. Furthermore, the PTMs studied here change the internal mechanic connectivity between key distal sectors of the protein required for proper functioning in normal conditions.

The overall outcome of the combination of these factors is the malfunction of the glycosylated protein: in this picture, PTMs perturb the population of GRP94 structural ensembles that are presented to the other partners necessary to assemble an efficient folding machinery, and consequently of the kinetics/timing of the chaperone cycle^35–38,57,58^. The net result is a perturbation of the foldase function. In terms of PPI networks, this translates into a remodeling of protein connectivity and, given the centrality of GRP94, the final effect is amplified in proteome-wide dysfunction.

Our extensive atomistic simulations also shed light on the determinants of GRP94 response to ligands. Interestingly, PU-WS13 dramatically perturbs the dynamics of the protein variants with glycosylation at N62, while forming a stable complex with the protein, traits that can help explain the activity of this drug in cells with high levels of ^Glyc62^GRP94. In contrast, the effects of PU-WS12 are much less pronounced with the ligand unable to populate a stable configuration for the bound state, which correlates with its apparent inactivity.

A key aspect of our work is that the comparative analysis of the dynamics of different glycosylated variants of GRP94 supports an active key role for glycosylation in regulating the conformational dynamics of the active site, where ATP and ATP-competitive ligands bind. The ATP-lid of GRP94 has a distinctive propensity to adopt the open structure, thus presenting an energetic barrier to closure of the dimer into its active conformation ^15,43,53,59^. While lid closure is required for ATP hydrolysis and for the chaperone cycle to proceed, *N62*-glycans stabilize the lid in open states, determine additional barriers for the lid to reach the closed conformation, and disfavor the population of the foldase state of the chaperone: this is consistent with the experimental characterization of N62-glycosylation as a key factor for the switch of GRP94 to a templating protein^15^. It is worth noting here that the conformational dynamics of the lid is substantially asymmetric between the two protomers, a theme already observed in the regulation of the functions of various members of the Hsp90 family ^39,41,42,45,51,54,60–63^. The observed sugar-induced asymmetry in the conformations of this substructure provides further potential for the expansion of the ensembles of GRP94 conformations that can interact with different partners, supporting the role of the glycosylated chaperone as the hub for proteome-wide interaction perturbations.

Overall, we suggest a picture whereby PTMs modulate the dynamics of the ATP-state as well as the efficiency of the catalytic hydrolysis cycle of the nucleotide, potentially providing a convenient mechanism of GRP94 adaptation to different cell conditions, in particular to chronic cellular stress states^64–66^, inducing extensive protein covalent modifications^10,67,68^.

Finally, our data provide a structure-based rationale for the observed (in)sensitivity of ^Glyc62^GRP94 to some GRP94 inhibitors. These results have important reach and implications in chemical biology and drug design as they suggest the possibility to leverage the structural and dynamic models generated here as templates to guide the selection and development of small molecules able to selectively act on cells where GRP94 aberrant glycosylation is a factor in determining pathologic states. More specifically, these small molecules could function by specifically engaging PTM-induced dynamic states, representing an important expansion of the molecular diversity space of chaperone targeting drugs ^20,56,69^. An additional advantage is the opportunity to selectively interfere with the disease-associated forms of the protein, leaving unmodified forms of normal cells untouched. Together with the development of other isoform specific ligands, this could decisively enrich our arsenal of chemical tools to clinically target and study the complex mechanisms of chaperones in their endogenous context. In conclusion, while based on the case of GRP94, our models and considerations are fully general, and readily transferable to other targets and contexts for cancers and other diseases.

### Significance

Aberrant *N*-glycosylation remodels GRP94 conformational dynamics, and, in turn, its selection of, and interaction with, protein partners. The final result is a remodeling of the connectivity within the PPI networks of which GRP94 is a central hub. The outcome at the cell level is that modified protein interaction pathways determine aberrant disease phenotypes. From these results, we propose a general model that can establish a link between the fundamental physicochemical basis of protein conformational behavior and the cellular functions it is involved in. The impact of protein modifications reflects the extent of the conformational dynamic perturbation and population shift between the functional and the altered forms of the protein. When considered in large systems, with multiple complexes that need to form and/or disrupt in a highly regulated, orderly manner with numerous players involved, these modifications can have significantly amplified effects on physiological signaling and deregulation in disease. Protein modifications and the variation of protein assembly mechanisms can thus reverberate in wide-spread proteome malfunctions determined by altered PPIs, which ultimately causes disease states.

The overall approach we present here may represent an effective means of exploiting the functional dynamics of different forms of GRP94, modulated by covalent modifications, in the selection and design of drugs that are specific and selective for a given form of the protein. Such drugs would intervene by specifically addressing PTM-induced dynamic states and by selectively interfering with the disease-associated form of the protein, leaving the unmodified forms of normal cells untouched. Together with the development of isoform specific ligands, this could decisively enrich our arsenal of chemical tools to clinically target and study the complex mechanisms of chaperones in their endogenous context. While based on the case of GRP94, our models and considerations are fully general, and readily transferable to other targets and contexts for cancers and other diseases.

## STAR+METHODS

Detailed methods are provided in the online version of this paper and include the following:

- KEY RESOURCES TABLE
- RESOURCE AVAILABILITY

- Lead Contact
- Material Availability
- Data and Code Availability
- EXPERIMENTAL MODEL AND SUBJECT DETAILS

- Human Cell Lines
- METHOD DETAILS

- Reagents
- Fluorescence polarization measurements
- SDS-PAGE and Immunoblotting
- Western blot analysis of signaling and viability
- GRP94 conformation detection by immunoprecipitation
- Native Gel Electrophoresis and GRP94 epichaperome detection
- ATP based cell viability assessment.
- Chemical synthesis and information
- Computational Methods
- System preparation
- Forcefield parameters for molecular dynamics simulations and parametrization of PU-WS12 and PU-WS13
- Molecular dynamics simulations
- Analysis of MD trajectories
- QUANTIFICATION AND STATISTICAL ANALYSIS

## SUPPLEMENTAL INFORMATION

Supplemental Information includes 11 figures and can be found with this article online at the journal website

## ACKNOWLEDGMENTS

This work is supported in part by the US National Institutes of Health (NIH) (R56 AG061869, P01 CA186866, P30 CA08748) for Gabriela Chiosis. Giorgio Colombo acknowledges funding from Fondazione AIRC (Associazione Italiana Ricerca Sul Cancro) grant IG 20019 and from PRIN (grant 20209KYCH9); The research leading to these results has received funding from AIRC under IG 2022 -ID 27139 project, PI Giorgio Colombo.

## AUTHOR CONTRIBUTIONS

M.C., P.Y., A.R., C.S.D., P.P. acquisition of data, analysis and interpretation of data. M.C., C.S.D., E.M., G.Chiosis, G. Colombo and drafting and revising the article. G. Chiosis, E.M., and G.Colombo conception and design of the study and drafting or revising the article.

## DECLARATION OF INTERESTS

G.Chiosis, A.R., C.S.D. and P.Y. are inventors on patents covering PU-WS13 and associated composition of matter. G.Chiosis. is a founder of Samus Therapeutics and a member of its board of directors.

## STAR+METHODS

### KEY RESOURCE TABLE

### RESOURCE AVAILABILITY

#### Lead Contact

Further information and requests for resources and reagents should be directed to the lead contacts, Dr. Giorgio Colombo and Dr. Elisabetta Moroni. For chemical compounds, requests should be addressed to Dr. Gabriela Chiosis.

#### Materials Availability

All unique reagents generated in this study are available from the Lead Contact with a completed Materials Transfer Agreement.

#### Data and Code Availability

The input files, topologies and restart files are provided as a zipped file. Any additional information required to reanalyze the data reported in this work is available at the link in the Key resource table.

### EXPERIMENTAL MODEL AND SUBJECT DETAILS

#### Human Cell Lines

The breast cancer cell lines MDA-MB-468 (HTB-132) and SKBr3 (HTB-30) were obtained from the American Type Culture Collection (ATCC) and were cultured in DMEM and McCoy’s 5A mediums supplemented with 10% FBS, 1% Glutamax and 1% penicillin and streptomycin (Pen/Strep), respectively. Both cell lines were incubated in the humidified cell incubators with 5% CO2 at 37 °C. Cells were authenticated using short tandem repeat profiling and tested for mycoplasma before and after use.

### METHOD DETAILS

#### Reagents

^Glyc62^GRP94 active ligands (PU-WS13, PU-SO33, PU-HJP149, PU-HJP110) and inactive ligands (BnIm, PU-WS12, PU-SO116, PU-HJP92, PU-HJP36) were synthesized using previously reported protocols ^15,24,30^. Briefly, for PU-WS13, a CuI-catalyzed coupling of 8-mercaptoadenine with 3,5-dichloroiodobenzene at 110°C resulted in 8-(3,5-dichloro-phenylsulfanyl)adenine in 72% yield, which was heated with 3-(*tert*butoxycarbonyl-isopropyl-amino)-propyl tosylate in DMF at 80°C under nitrogen protection for 30 min. The boc-deprotection of resulting *tert*-butyl(3-(6-amino-8-((3,5-dichlorophenyl)thio)-9*H*-purin-9-yl)propyl)(isopropyl) carbamate from the previous step with TFA gave crude material which upon purification by preparatory TLC with CH_2_Cl_2_:MeOH-NH_3_ (7N) at 20:1 gave **PU-WS13** in 45% yield. ^1^H NMR (600 MHz, CDCl_3_/CD_3_OD): δ 8.21 (s, 1H), 7.26 (t, *J* = 1.7 Hz, 1H), 7.24 (d, *J* = 1.9 Hz, 2H), 4.23 (t, *J* = 6.9 Hz, 2H), 2.63 (septet, *J* = 6.2 Hz, 1H), 2.46 (t, *J* = 6.4 Hz, 2H), 1.89 (quintet, *J* = 6.9 Hz, 2H), 0.97 (d, *J* = 6.3 Hz, 6H); ^13^C NMR (150 MHz, CDCl3): δ 154.7, 153.1, 151.3, 144.1, 135.9, 133.7, 128.8, 128.7, 119.7, 48.6, 43.4, 41.6, 29.8, 22.4; HRMS (ESI): m/z [M+H]^+^ calcd for C_17_H_21_Cl_2_N_6_S 411.0925, found 411.0917. For PU-WS12, CuI-catalyzed coupling of 8-mercaptoadenine with 1-iodo-2,4-dichlorobenzene at 110°C resulted in 8-((2,4-dichlorophenyl)thio)-9*H*-purin-6-amine in 87% yield, which was heated with 3-(*tert*butoxycarbonyl-isopropyl-amino)-propyl tosylate in DMF at 80°C under nitrogen protection for 30 min. The boc-deprotection of resulting *tert*-butyl (3-(6-amino-8-((2,4-dichlorophenyl)thio)-9*H*-purin-9-yl)propyl)(isopropyl)carbamate from the previous step with TFA gave crude material which upon purification by preparatory TLC with CH_2_Cl_2_:MeOH-NH_3_ (7N) at 20:1 yielded **PU-WS12**(35% yield). ^1^H NMR (600 MHz, CDCl_3_/CD_3_OD): δ 8.23 (s, 1H), 7.55 (s, 1H), 7.38 (d, *J* = 8.3 Hz, 1H), 7.30 (d, *J* = 8.3 Hz, 1H), 4.34 (t, *J* = 6.7 Hz, 2H), 2.92 (septet, *J* = 5.8 Hz, 1H), 2.69 (t, *J* = 6.4 Hz, 2H), 2.13 (quintet, *J* = 6.5 Hz, 2H), 1.16 (d, *J*= 5.7 Hz, 6H); HRMS (ESI): m/z [M+H]^+^ calcd for C_17_H_21_Cl_2_N_6_S 411.0925, found 411.0907. Synthesis of PU-SO33 includes cyclocondensation of 2,4,5,6-tetraaminopyrimidine sulfate with CS_2_ in ethanol to obtain 2,6-diamino-9*H*-purine-8-thiol in quantitative yield. Next, copper-catalyzed coupling of 2,6-diamino-9*H*-purine-8-thiol with 3,5-dichloroiodobenzene resulted in 65% yield of 8-((3,5-dichlorophenyl)thio)-9*H*-purine-2,6-diamine. Transformation of C2-amino group of 8-((3,5-dichlorophenyl)thio)-9*H*-purine-2,6-diamine to Cl was achieved with SbCl_3_ and *t*-BuONO in 10:1 mixture of DCE and DMSO at 80°C resulting in 2-chloro-8-((3,5-dichlorophenyl)thio)-9*H*-purin-6-amine. The sequential reaction of 2-chloro-8-((3,5-dichlorophenyl)thio)-9*H*-purin-6-amine with 1,3-dibromopropane, followed by treating the bromo derivative from the previous step with excess isopropylamine resulted in crude **PU-SO33**. The crude residue was purified by preparatory TLC (CH_2_Cl_2_-MeOH-NH_3_ (7N), 20:1) to give 60% yield of **PU-SO33**. ^1^H NMR (500 MHz, CDCl_3_) δ 7.33 – 7.31 (m, 1H), 7.30 – 7.28 (m, 2H), 5.75 (br s, 2H), 4.29 (t, *J* = 6.9 Hz, 2H), 2.78 – 2.70 (m, 1H), 2.56 (t, *J* = 6.6 Hz, 2H), 1.99 – 1.91 (m, 2H), 1.05 (d, *J* = 6.2 Hz, 6H); ^13^C NMR (150 MHz, CDCl_3_) δ 155.0, 154.5, 152.7, 144.6, 135.9, 134.0, 128.7, 128.5, 118.9, 48.9, 43.6, 42.0, 30.1, 22.7; MS m/z 444.92 [M+H]^+^. For methyl 2-(2-(1-benzyl-1*H*-imidazol-2-yl)ethyl)-3-chloro-4,6-dihydroxybenzoate (**BnIm**), the reaction of methyl 2,4-dihydroxy-6-methylbenzoate with *tert*-butyldimethylsilyl chloride in the presence of NaH provided TBS-protected methyl benzoate derivative, which was treated with calcium hypochlorite in 10% AcOH/H_2_O to achieve chlorination. The resulting compound from the previous step reacted with ally bromide and lithium diisopropylamide at −78°C in THF to obtain methyl 2-(but-3-en-1-yl)-4,6-bis((*fe/τ*-butyldimethylsilyl)oxy)-3-chlorobenzoate in 45% yield. Next, the oxidation of 2-(but-3-en-1-yl)-4,6-bis((*ferf*-butyldimethylsilyl)oxy)-3-chlorobenzoate with ozone and the reaction of resulting aldehyde compound with benzylamine and glyoxal, followed by TBS-deprotection with TBAF provided crude BnIm compound. The purification was achieved using column chromatography (CH_2_Cl_2_-MeOH, 20:1) which provided 22% yield of **BnIm**. ^1^H NMR (600 MHz, CDCl_3_) δ 11.35 (s, 2H), 7.40 – 7.36 (m, 2H), 7.35 – 7.32 (m, 1H), 7.10 – 7.07 (m, 3H), 6.90 (d, *J* = 1.3 Hz, 1H), 6.52 (s, 1H), 5.14 (s, 2H), 3.86 (s, 3H), 3.58 – 3.54 (m, 2H), 3.01 – 2.97 (m, 2H); ^13^C NMR (150 MHz, CDCl3) δ 170.8, 162.8, 158.1, 147.6, 141.5, 136.1, 129.0, 128.1, 126.6, 120.1, 114.9, 105.8, 102.7, 52.5, 49.5, 30.8, 26.0; HRMS (ESI): m/z [M+H]^+^ calcd for C_20_H_20_ClN_2_O_4_ 387.1112, found 387.1116. For the sysnthesis and characterization of PU-HJP149, PU-HJP110, PU-SO116 and PU-HJP92 and PU-HJP36 refer to Methods, **Chemical synthesis and information.**

#### Fluorescence polarization (FP) measurements

The FP competition assays were performed on an Analyst GT instrument (Molecular Devices, Sunnyvale, CA) and carried out in black 96-well micro-plates (Corning, no. 3650) in a total volume of 100 μL in each well. A stock of 10 μM cy3B-GM ^70^ was prepared in DMSO and diluted with Felts buffer (20 mM Hepes (K), pH 7.3, 50 mM KCl, 2 mM DTT, 5 mM MgCl_2_, 20 mM Na_2_MoO_4_ and 0.01% NP40 with 0.1 mg/mL BGG). To each well was added 6 nM cy3B-GM, 10 nM recombinant GRP94 or HSP90α protein and tested inhibitor (initial stock in DMSO) in a final volume of 100 μL Felts buffer. Compounds were added in triplicate wells. For each assay, background wells (buffer only), tracer controls (free, cy3B-GM only) and bound controls (cy3B-GM in the presence of protein) were included on each assay plate. The assay plate was incubated on a shaker at 4 °C for 24 h, and the FP values (in mP) were measured. The fraction of cy3B-GM bound GRP94 was correlated to the mP value and plotted against values of competitor concentrations. The inhibitor concentration at which 50% of bound cy3B-GM was displaced was obtained by fitting the data. An excitation filter at 530 nm and an emission filter at 580 nm were used with a dichroic mirror of 561 nm. All of the experimental data were analyzed using SOFTmax Pro 4.3.1 and plotted using Prism 6.0 (GraphPad Software Inc., San Diego, CA). Binding affinity values were given as relative binding affinity values (EC_50_, concentration at which 50% of fluorescent ligand was competed off by compound).

#### SDS-PAGE and Immunoblotting

Cells were either treated with DMSO (vehicle) or indicated compounds and lysed in RIPA buffer (50 mM Tris-HCl, pH 7.5, 150 mM NaCl, 0.1% sodium deoxycholate and 0.5% NP40) supplemented with cocktail protease inhibitors (Roche) to produce whole-cell lysates (WCLs). Membrane fraction was prepared by ProteoExtract Subcellular Proteome Extraction Kit (Millipore, 539790) according to the manufacturer’s instructions. Protein concentrations were determined using the BCA kit (Pierce). Ten to fifty micrograms of total protein were examined by immunoblotting with indicated antibodies. GRP94 antibody (SPA-850, 1:3,000) from Enzo, HER2 antibody (28-0004, 1:2,000) from Invitrogen, EGFR (#4267, 1:1,000), p-ERK1/2 (#4377, 1:2,000) and total ERK1/2 (#4695, 1:3,000) antibodies from Cell Signaling Technology were used. The blots were washed with TBS/0.1% Tween20 and incubated with appropriate HRP-conjugated secondary antibodies (1:3,000). Chemiluminescent signal was detected with Enhanced Chemiluminescence Detection System (GE Healthcare) according to the manufacturer’s instructions.

#### Western blot analysis of signaling and viability

MDA-MB-468 cells were plated in 6-well plates at 3 x 10^6^ cells per plate and treated next day with the indicated concentrations of PU-WS12, PUWS13 or vehicle. The cells were collected after 24 h of treatment and protein extracts were prepared in RIPA buffer. Twenty micrograms of total protein were examined by immunoblotting with cleaved PARP (5625, 1:1,000), phospho-ERK (T202/Y204) (4377, 1:3,000) and ERK (4695, 1:5,000) antibodies purchased from Cell Signaling Technology. The blots were processed as described in the immunoblotting section.

#### GRP94 conformation detection by immunoprecipitation (IP)

*In cell IP:* MDA-MB-468 cells or SkBr3 cells were plated in 10 cm plates at 3 x 10^6^ cells per plate and treated next day with the indicated compounds at indicated concentrations. At the end of the treatment cells were washed with cold PBS and harvested. Protein extracts were prepared in the RIPA buffer. 100 μg whole cell lysates were incubated with 2 μg either 9G10 (SPA-850, Enzo) or G4420 (Sigma) antibody for 3 hr at 4°C, followed by incubation with Protein A/G agarose beads (Roche) for another 2 hr at 4°C. The beads were washed with cold Felts buffer three times, eluted by SDS loading buffer and subjected to immunoblotting as described above. *In lysate IP:* MDA-MB-468 cells were washed with cold PBS, harvested by centrifugation and lysed in RIPA buffer. 200 μg cell lysates were diluted to 100 μL with Felts buffer and incubated with 1 μM PU-WS13 or PU-WS12 for 2 hr at 4°C. 2 μg 9G10 antibody (SPA-850, Enzo) was added to the samples and incubated at 4°C overnight, followed by incubation with Protein G agarose beads for another 2 hr at 4°C. The beads were washed with cold Felts buffer three times, eluted by SDS loading buffer and subjected to immunoblotting as described above.

#### Native Gel Electrophoresis and GRP94 epichaperome detection

Cells were lysed in the RIPA buffers (WCL) and diluted with Felts buffer (20 mM Hepes pH 7.3, 50 mM KCl, 2 mM DTT, 5 mM MgCl_2_, 20 mM Na_2_MoO_4_ and 0.01 % NP40 with 0.1 mgmL^-1^ BGG). Twenty-five to one hundred μg of protein were loaded onto 5% native gel and resolved at 4°C. The gels were stained with Bio-Safe Coomassie stain (Biorad, 1610786), or soaked in Tris-Glycine-SDS running buffer for 15 min prior to gel transfer for 75 min at 100V and immunoblotting for GRP94 using the 9G10 antibody as described above.

#### ATP based cell viability assessment

Cell viability was assessed using CellTiter-Glo luminescent Cell Viability Assay (Promega) according to the manufacturer’s instructions. The method determines the number of viable cells in culture based on quantification of ATP amount, which signals the presence of metabolically active cells, and was performed as previously reported (Yan 2020 cell reports). Briefly, MDA-MB-468 cells were plated in clear bottom black 96-well plates (Corning) at 6,000 cells per well and treated with serial dilutions of the indicated compounds for 72 h. After incubation, 100 μL of CellTiter-Glo reagent was added to each well. Plates were incubated with the reagent for 10 min at room temperature and the luminescence signal was measured using Analyst GT microplate reader (Molecular Devices). IC50 values for each compound were determined using Prism GraphPad.

#### Chemical synthesis and information

##### Chemistry: General experimental

###### Chemistry general considerations

All commercial chemicals and solvents were reagent grade and used without further purification. The identity and purity of each product was characterized by MS, HPLC, TLC, and NMR. ^1^H/^13^C NMR spectra were recorded on either a Bruker 400, 500 or 600 MHz instrument. Chemical shifts are reported in δ values in ppm downfield from TMS as the internal standard. ^1^H data are reported as follows: chemical shift, multiplicity (s = singlet, d = doublet, t = triplet, q =quartet, br = broad, m = multiplet), coupling constant (Hz), integration. ^13^C chemical shifts are reported in δ values in ppm downfield from TMS as the internal standard. Low resolution mass spectra were obtained on Waters Acquity Ultra Performance LC with electrospray ionization and SQ detector. Purity of target compounds has been determined to be >95% by LC/MS on a Waters Autopurification system with PDA, MicroMass ZQ and ELSD detector and a reversed phase column (Waters X-Bridge C18, 4.6 x 150 mm, 5 μm) eluted with water/acetonitrile gradients, containing 0.1% TFA. Column chromatography was performed using 230-400 mesh silica gel. Analytical thin layer chromatography was performed on 250 μM silica gel F254 plates. Preparative thin layer chromatography was performed on 1000 μM silica gel F254 plates. Flash chromatography was performed using CombiFlash^®^R_f_ instrument. Compounds **1**, **7**, **13**, and **16** were synthesized as reported previously ^15,30,71,72^.

##### Synthetic scheme of PU-HJP36

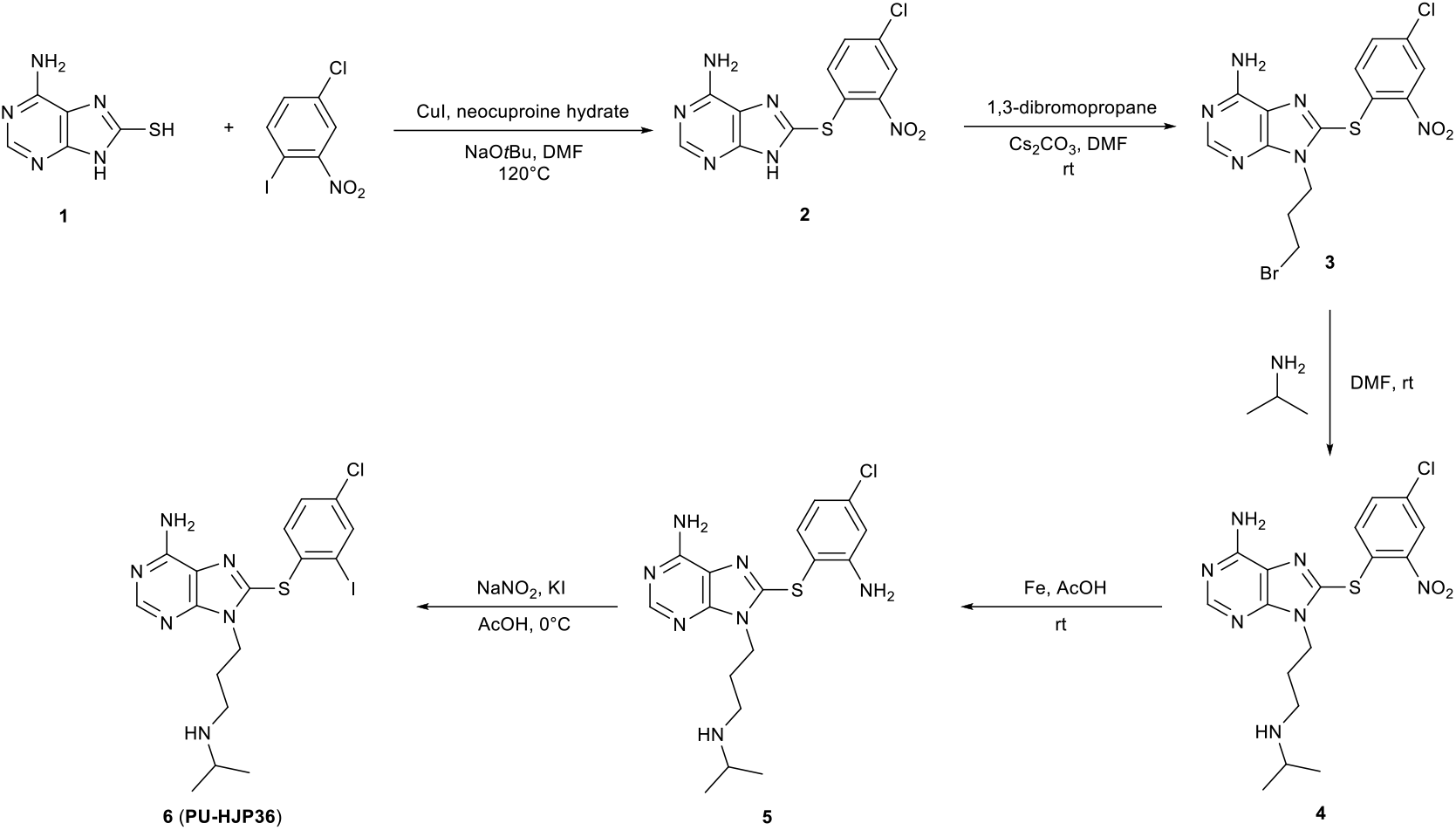

###### Step 1: Preparation of 8-((4-Chloro-2-nitrophenyl)thio)-9H-purin-6-amine (2)

8-Mercaptoadenine (**1**, 1.6 g, 8.8 mmol), neocuproine hydrate (540 mg, 2.6 mmol), Cul (330 mg, 1.73 mmol), NaO*t*Bu (1.7 g, 17.38 mmol), 4-chloro-1-iodo-2-nitrobenzene (3.2 g, 11.2 mmol), and anhydrous DMF (25 mL) were taken in a round bottom flask flushed with nitrogen. The flask was sealed with Teflon tape, heated at 120 °C, and magnetically stirred for 20 h under nitrogen. Solvent was removed under reduced pressure and the resulting residue was purified using flash chromatography (CH2Cl2:MeOH, 10:0 to 10:1) to provide **2**in 95 % yield (2.7 g). MS (ESI): m/z 322.8 [M + H]^+^.

###### Step 2 and 3: Preparation of 9-(3-Bromopropyl)-8-((4-chloro-2-nitrophenyl)thio)-9H-purin-6-amine (3) and 8-((4-chloro-2-nitrophenyl)thio)-9-(3-(isopropylamino)propyl)-9H-purin-6-amine (4)

Compound **2**(2 g, 6.211 mmol) was dissolved in DMF (20 mL) and Cs_2_CO_3_ (2.43 g, 7.45 mmol) and 1,3-dibromopropane (3.15 mL, 31.05 mmol) were added and the mixture was stirred under nitrogen at room temperature for 2 h. After completion of reaction, Cs_2_CO_3_ was filtered off and the residue was washed with DMF (3 mL). The filtrate containing crude compound **3** was taken into an RBF and directly reacted with isopropylamine (0.31 mmol) overnight. Following solvent removal, the crude material was purified by column chromatography (CH_2_Cl_2_:CH_3_OH-NH_3_ (7N), 100:1 to 20:1) to afford desired product **4** in 23% yield (600 mg). ^1^H NMR (600 MHz, CDCl3/CD3OD) δ 8.35 (s, 1H), 8.30 (d, *J*= 2.2 Hz, 1H), 7.43 (dd, *J* = 8.7, 2.2 Hz, 1H), 6.81 (d, *J*= 8.7 Hz, 1H), 4.29 (t, *J* = 6.9 Hz, 2H), 2.67-2.73 (m, 1H), 2.52 (t, *J* = 6.8 Hz, 2H), 1.93-1.98 (m, 2H), 1.03 (d, *J* = 6.3 Hz, 6H); ^13^C NMR (150 MHz, CDCl3/CD3OD) δ 155.3, 153.9, 151.3, 145.8, 142.0, 134.5, 133.3, 131.9, 129.7, 126.2, 120.4, 48.7, 43.3, 41.9, 30.1, 22.3; HRMS (ESI) m/z [M+H]^+^ calcd. for C17H21ClN7OS, 422.1166; found 422.1170.

###### Step 4: Preparation of 8-((2-Amino-4-chlorophenyl)thio)-9-(3-(isopropylamino)propyl)-9H-purin-6-amine (5)

A mixture of **4**(300 mg, 0.711 mmol) and iron powder (150 mg, 2.488 mmol) in acetic acid (5 mL) was stirred at room temperature for 4 hrs. On completion, the reaction was neutralized by adding solid Na_2_CO_3_ at 0 °C and washed with EtOAc (75 ml X 3). Following drying the organic layer over MgSO_4_ and solvent removal, the crude material was purified by column chromatography (CH_2_Cl_2_:CH_3_OH-NH_3_ (7N), 50:1 to 15:1) to afford **5**in 93% yield (260 mg). ^1^H NMR (600 MHz, CDCl3/CD3OD) δ 8.16 (s, 1H), 7.39 (d, *J* = 8.3 Hz, 1H), 6.84 (d, *J* = 2.2 Hz, 1H), 6.73 (dd, *J* = 8.3, 2.2 Hz, 1H), 4.36 (t, *J* = 6.7 Hz, 2H), 3.40-3.41 (m, 1H), 2.83 (t, *J* = 6.8 Hz, 2H), 2.26-2.32 (m, 2H), 1.34 (d, *J* = 6.5 Hz, 6H); ^13^C NMR (150 MHz, CDCl_3_/CD_3_OD) δ 154.2, 151.9, 151.4, 150.5, 147.3, 138.3, 138.1, 118.9, 115.7, 107.0, 74.4, 50.6, 42.1, 40.4, 31.1, 19.8; HRMS (ESI) m/z [M+H]^+^ calcd. for C_17_H_23_ClN_7_S, 392.1424; found 392.1419.

###### Step 5: 8-((4-Chloro-2-iodophenyl)thio)-9-(3-(isopropylamino)propyl)-9H-purin-6-amine (6, PU-HJP36)

A mixture of **5**(260 mg, 0.665 mmol), NaNO_2_ (57 mg, 0.831 mmol) and potassium iodide (221 mg, 1.3 mmol) in acetic acid (5 mL) was stirred at 0°C for 15 min. On completion, water (10 mL) was added, and the reaction was neutralized by adding solid Na_2_CO_3_ at 0°C and further washed with EtOAc (75 ml X 3). Following drying over MgSO_4_ and solvent removal, the crude material was purified by column chromatography (CH_2_Cl_2_:CH_3_OH-NH_3_ (7N), 80:1 to 20:1) to afford **PU-HJP36**(**6**) in 79% yield (260 mg). ^1^H NMR (600 MHz, CDCl_3_) δ 8.35 (s, 1H), 7.87 (d, *J* = 2.2 Hz, 1H), 7.23 (dd, *J* = 8.5, 2.2 Hz, 1H), 7.07 (d, *J* = 2.2 Hz, 1H), 5.87 (br s, 2H), 4.30 (t, *J* = 6.9 Hz, 2H), 2.68-2.72 (m, 1H), 2.56 (t, *J* = 6.8 Hz, 2H), 1.95-1.99 (m, 2H), 1.03 (d, *J* = 6.2 Hz, 6H); ^13^C NMR (150 MHz, CDCl_3_) δ 154.7, 153.3, 151.6, 144.8, 139.3, 135.9, 134.1, 131.1, 129.5, 120.4, 99.8, 48.7, 43.8, 41.9, 30.3, 22.9; HRMS (ESI) m/z [M+H]^+^ calcd. for C_17_H_21_IClN_6_S, 503.0282; found 503.0277.

##### Synthetic scheme of PU-SO116

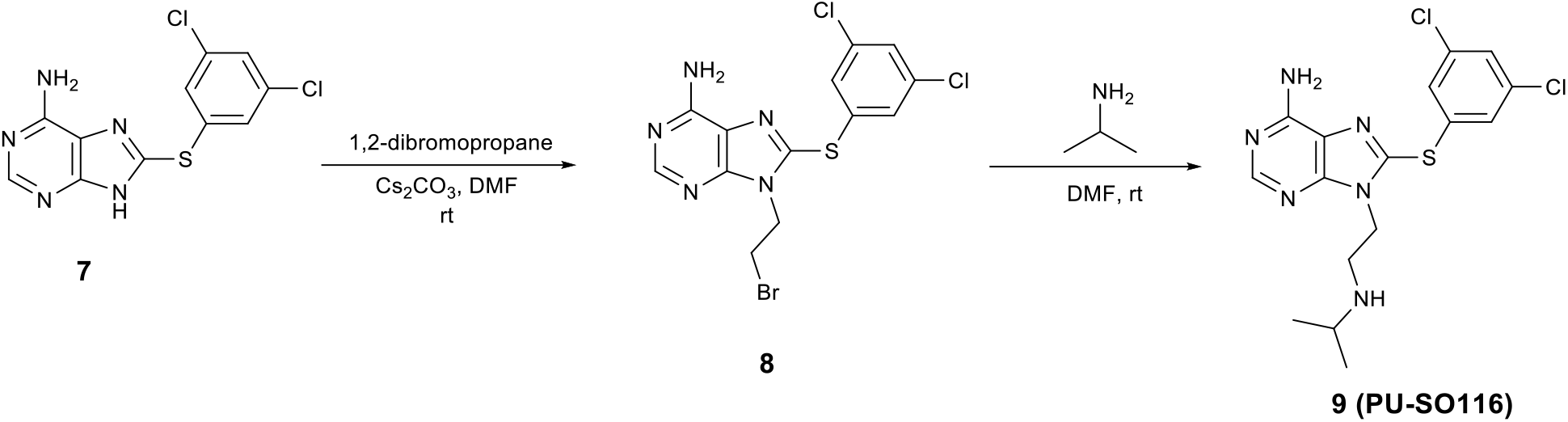

###### Step 1: 9-(2-Bromoethyl)-8-((3,5-dichlorophenyl)thio)-9H-purin-6-amine (8)

To a solution of **7** (200 mg, 0.640 mmol) in DMF (5 mL) was added Cs2CO3 (335 mg, 1.025 mmol) and 1,2-dibromoethane (277 μL, 3.2 mmol) and the mixture was stirred under nitrogen at rt for 4 h. Solvent was removed under reduced pressure and the resulting residue was chromatographed (CH_2_Cl_2_:MeOH:AcOH, 20:1:0.5) to afford **8** in 38 % yield (102 mg). NMR (400 MHz, CDCl_3_) δ 8.34 (s, 1H), 7.28 (s, 2H), 7.25 (s, 1H), 6.54 (s, 2H), 4.66 (t, *J* = 6.5 Hz, 2H), 3.74 (t, *J* = 6.5 Hz, 2H); MS (ESI): m/z 420.2 [M+H]^+^.

###### Step 2: 8-((3,5-Dichlorophenyl)thio)-9-(2-(isopropylamino)ethyl)-9H-purin-6-amine (9, PU-SO116)

To **8**(20 mg, 0.047 mmol) in dry DMF (2 mL) was added isopropylamine (121 μL, 1.41 mmol) and the reaction mixture was stirred at rt for 72 hours. Solvent was removed under reduced pressure and the residue was purified by preparatory TLC (CH_2_Cl_2_:MeOH-NH_3_ (7N), 20: 1) to afford 12.6 mg (68%) of **PU-SO116**. ^1^H NMR (600 MHz, CDCl_3_): δ 8.36 (s, 1H), 7.26-7.28 (m, 3H), 5.82 (br s, 2H), 4.34 (t, *J* = 6.5 Hz, 2H), 2.97 (t, *J* = 6.5 Hz, 2H), 2.74-2.76 (m, 1H), 0.96 (d, *J* = 6.2 Hz, 6H). ^13^C NMR (150 MHz, CDCl_3_): δ 154.8, 153.5, 151.5, 144.0, 135.7, 135.3, 128.1, 127.8, 120.4, 48.5, 46.1,44.5, 22.8. HRMS (ESI) m/z [M+H]^+^ calcd. for C_16_H_19_N_6_SCl_2_, 397.0769; found 397.0765.

##### Synthetic scheme of PU-HJP149

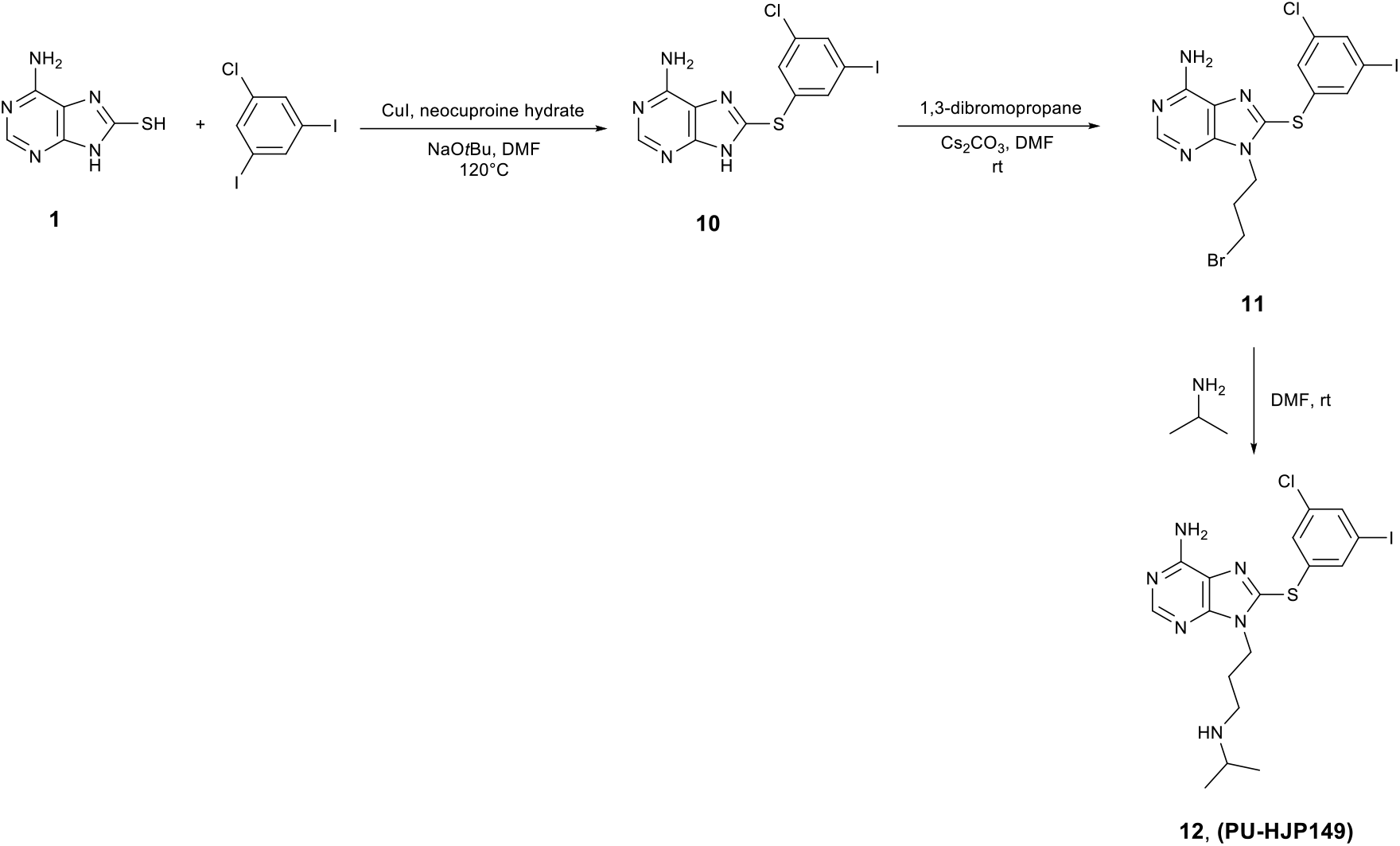

###### Step 1: 8-((3-chloro-5-iodophenyl)thio)-9H-purin-6-amine (10)

8-Mercaptoadenine (**1**, 1 g, 5.988 mmol), neocuproine hydrate (0.25 g, 1.1976 mmol), Cul (0.23 g, 1.1976 mmol), NaO*t*Bu (1.45 g, 11.976 mmol), 1-chloro-3,5-diiodobenzene (3.3 g, 8.982 mmol), and anhydrous DMF (25 mL) were taken in a round bottom flask flushed with nitrogen. The flask was sealed with Teflon tape, heated at 120 °C, and magnetically stirred for 20 h under nitrogen. Solvent was removed under reduced pressure and the resulting residue was chromatographed (CH_2_Cl_2_:MeOH: 10:1) to provide **10** in 67 % yield (1.6 g). ^1^H NMR (500 MHz, DMSO-*d_6_*) δ 13.47 (br s, 1H), 8.13 (s, 1H), 7.81 (s, 1H), 7.72 (s, 1H), 7.50 (s, 1H), 7.37 (br s, 2H); MS (ESI): m/z 403.7 [M + H]^+^.

###### Step 2 and 3: 9-(3-Bromopropyl)-8-((3-chloro-5-iodophenyl)thio)-9H-purin-6-amine (11) and 8-((3-chloro-5-iodophenyl)thio)-9-(3-(isopropylamino)propyl)-9H-purin-6-amine (12, PU-HJP149)

To a solution of **10**(500 mg, 1.24 mmol) in DMF (15 mL) was added Cs_2_CO_3_ (485 mg, 1.48 mmol) and 1,3-dibromopropane (630 μL, 6.2 mmol). The reaction mixture was stirred under nitrogen at room temperature for 2 h. On completion, the solid was filtered off and washed with DMF (4 mL). The filtrate containing crude compound **11** was taken into an RBF and directly reacted with isopropylamine (750 μL) at room temperature overnight. Solvent was removed under reduced pressure and the resulting residue was purified by preparative TLC (CH_2_Cl_2_:CH_3_OH-NH_3_ (7N), 20: 1 or 60: 1) to afford **PU-HJP149**(**12**) in 20% yield (125 mg). ^1^H NMR (600 MHz, CDCl_3_) δ 8.34 (s, 1H), 7.62 (dt, *J* = 7.4 Hz, 1.4 Hz, 2H), 7.34 (t, *J* = 1.7 Hz, 1H), 6.13 (br s, 2H), 4.32 (t, *J* = 7.0 Hz, 2H), 2.67-2.73 (m, 1H), 2.55 (t, *J* = 6.8 Hz, 2H), 1.92-1.98 (m, 2H), 1.03 (d, *J* = 6.2 Hz, 6H); ^13^C NMR (150 MHz, CDCl_3_) δ 154.8, 153.4, 151.6, 143.6, 136.7, 136.6, 135.7, 134.7, 129.3, 120.2, 94.4, 48.7, 43.8, 41.8, 30.3, 22.9; HRMS (ESI) m/z [M+H]+ calcd. for C_17_H_21_IClN_6_S, 503.0282; found 503.0260.

##### Synthetic scheme of PU-HJP92

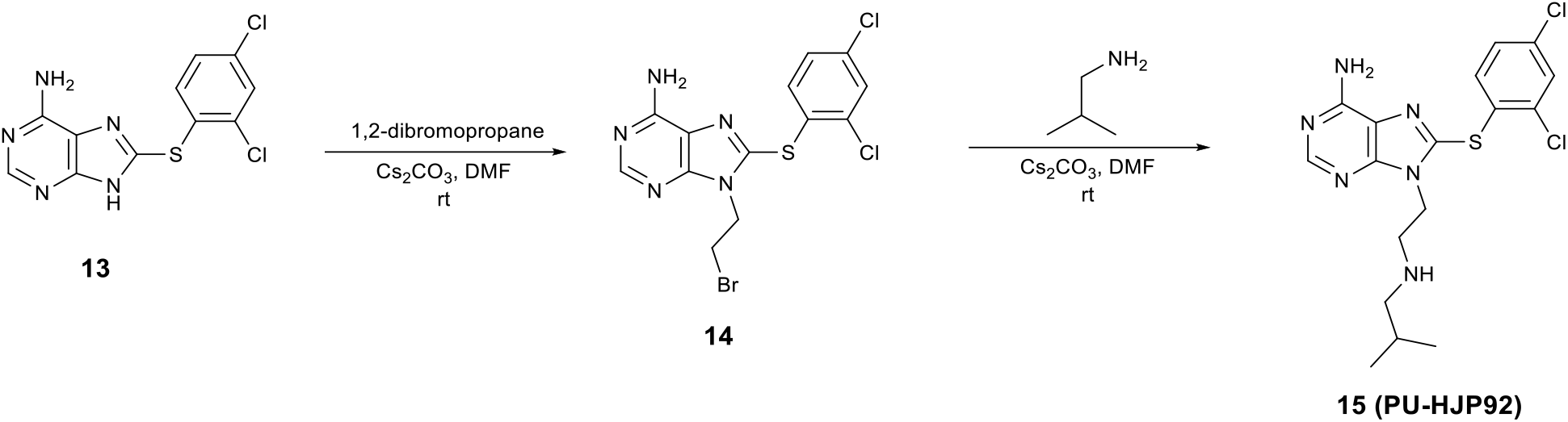

###### Step 1: 9-(2-bromoethyl)-8-((2,4-dichlorophenyl)thio)-9H-purin-6-amine (14)

A mixture of **13** (400 mg, 1.28 mmol), Cs_2_CO_3_ (630 mg, 1.92 mmol), and 1,2-dibromopropane (1.21 g, 0.55 mL, 6.43 mmol) in DMF (10 mL) under nitrogen protection was stirred at rt for 3 h. Following solvent removal, the crude material was purified by preparatory TLC (CH_2_Cl_2_:CH_3_OH:AcOH, 20:1:0.1) to afford **14** in 36% yield (190 mg). ^1^H NMR (500 MHz, CDCl_3_/CD_3_OD) δ 8.27 (s, 1H), 7.52 (d, *J* = 2.2 Hz, 1H), 7.36 (d, *J* = 8.5 Hz, 1H), 7.26 (dd, *J* = 8.4, 2.2 Hz, 1H), 4.68 (t, *J* = 6.5 Hz, 2H), 3.77 (t, *J* = 6.5 Hz, 2H); ^13^C NMR (125 MHz, CDCl_3_/CD_3_OD) δ 154.6, 153.1, 150.9, 145.0, 136.3, 135.7, 133.9, 130.3, 128.4, 128.2, 119.8, 45.0, 28.5; MS (ESI): m/z 417.9 [M + H]^+^.

###### Step 2: 8-((2,4-dichlorophenyl)thio)-9-(2-(isobutylamino)ethyl)-9H-purin-6-amine (15, PU-HJP92)

A mixture of **14** (9 mg, 0.022 mmol) and isobutylamine (53.5 μL, 0.54 mmol) in DMF (1 mL) under nitrogen protection was stirred at room temperature overnight. Following solvent removal, the crude material was purified by preparative TLC (CH_2_Cl_2_:CH_3_OH-NH_3_ (7N), 20:1) to afford **PU-HJP92** in 81% yield (7.3 mg). ^1^H NMR (600 MHz, CDCl_3_/CD_3_OD) δ 8.21 (s, 1H), 7.43 (d, *J* = 2.0 Hz, 1H), 7.22 (d, *J* = 8.0 Hz, 1H), 7.16 (dd, *J* = 8.5 Hz, 2.0 Hz, 1H), 4.31 (t, *J* = 5.9 Hz, 2H), 2.94 (t, *J* = 5.7 Hz, 2H), 2.34-2.37 (m, 2H), 1.62-1.66 (m, 1H), 0.80 (d, *J* = 6.5 Hz, 6H); ^13^C NMR (150 MHz, CDCl_3_/CD_3_OD) δ 154.5, 152.9, 151.2, 145.2, 136.1, 135.4, 133.7, 130.3, 130.1, 128.9, 128.2, 119.8, 57.3, 48.6, 43.7, 28.0, 20.4; HRMS (ESI) m/z [M+H]^+^ calcd. for C_17_H_21_Cl_2_N_6_S, 411.0925; found 411.0917.

##### Synthetic scheme of PU-HJP110

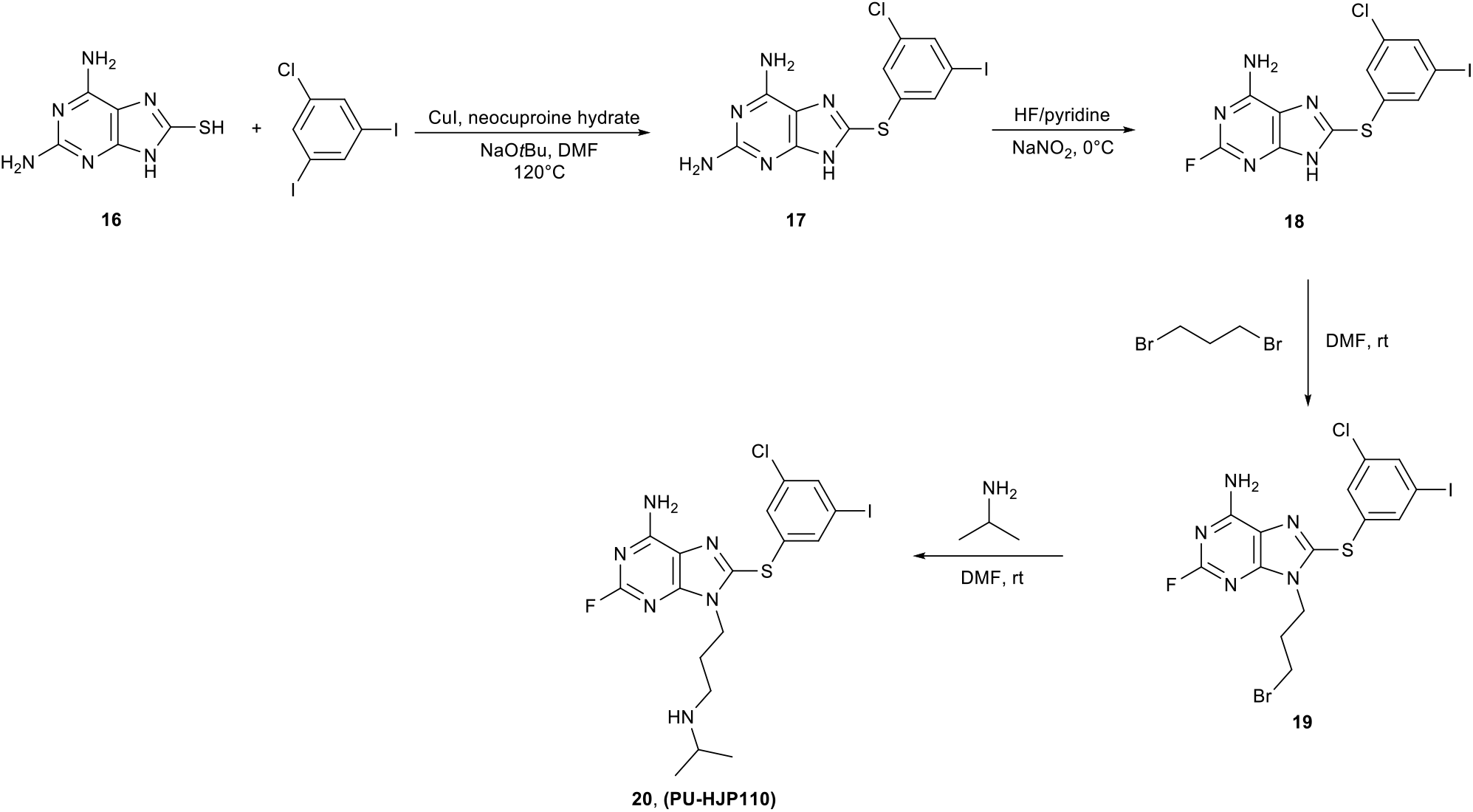

###### Step 1: 8-((3-chloro-5-iodophenyl)thio)-9H-purine-2,6-diamine (17)

Compound 16 (2 g, 10.989 mmol), 1-chloro-3,5-diiodobenzene (5 g, 13.736 mmol), neocuprine hydrate (0.46 g, 2.1978 mmol), Cul (0.42 g, 2.1978 mmol), NaOtBu (2.11 g, 21.978 mmol) and DMF (35 mL) were charged in a nitrogen protected dry vessel. The reaction vessel was sealed and placed in an oil bath (120°C) and stirred for 20 h. The reaction mixture was then cooled to room temperature and DMF was removed in vacuo. The crude material was purified by silica gel column chromatography (CH_2_Cl_2_:CH_3_OH, 10:0 to 10:1) to afford 3.3 g (71.8 %) of 17. MS (ESI): m/z 418.6 [M + H]^+^.

###### Step 2: 8-((3-chloro-5-iodophenyl)thio)-2-fluoro-9H-purin-6-amine (18)

To a cooled solution (0 °C) of **17** (1 g, 2.392 mmol) in HF/pyridine (1.5 mL) was slowly added NaNO_2_ (185 mg, 3.11 mmol). The resulted mixture was stirred at 0°C for 10 min and then at room temperature for 1.5 h. The reaction mixture was quenched by stirring with saturated CaCO3 solution (5 mL) for 1 h. The crude material was taken up in EtOAc, washed with water, and dried over anhydrous Na2SO4. Following solvent removal, the residue was purified using flash chromatography (CH_2_Cl_2_:CH_3_OH, 100:0 to 10:1) to afford **18**(100 mg, 11% yield). MS (ESI): m/z 421.8 [M + H]^+^.

###### Step 3: 9-(3-bromopropyl)-8-((3-chloro-5-iodophenyl)thio)-2-fluoro-9H-purin-6-amine (19)

A mixture of **18** (40 mg, 0.0951 mmol), Cs_2_CO_3_ (40.51 mg, 0.1236 mmol), and 1,3-dibromopropane (48.3 μL, 0.476 mmol) in DMF (2 mL) under nitrogen protection was stirred at room temperature for 2 h. Following solvent removal, the crude material was purified by preparatory TLC (CH_2_Cl_2_:CH_3_OH:AcOH, 40:1:0.1) to afford **19** in 45% yield (23 mg). ^1^H NMR (500 MHz, CDCl_3_) δ 7.64 (dt, *J* = 4.1, 1.5 Hz, 2H), 7.36 (t, *J* = 1.7 Hz, 1H), 6.42 (br s, 2H), 4.34 – 4.29 (m, 2H), 3.39 (t, *J* = 6.3 Hz, 2H), 2.36 – 2.29 (m, 2H); ^13^C NMR (125 MHz, CDCl_3_) δ 159.38 (d, *J* = 212.5 Hz), 156.61 (d, *J* = 20.2 Hz), 152.94 (d, *J* = 19.4 Hz), 143.71 (d, *J* = 2.8 Hz), 137.10, 136.90, 135.89, 134.19, 129.58, 118.46 (d, *J* = 3.8 Hz), 94.58, 42.82, 32.03, 29.18; MS m/z 541.8 [M + H]^+^.

###### Step 4: 8-((3-chloro-5-iodophenyl)thio)-2-fluoro-9-(3-(isopropylamino)propyl)-9H-purin-6-amine (20, HJP-VI-110)

To a solution of 19 (23 mg, 0.00423 mmol) in dry DMF (2 ml) was added isopropylamine (173 μL, 2.1194 mmol) and the reaction mixture was stirred at rt for 8 h. Then, the solvent was removed under reduced pressure and the crude product was purified by prep TLC (CH_2_Cl_2_:MeOH-NH_3_ (7N), 20:1) to afford 15.2 mg (69%) of 20 (HJP-V-110). ^1^H NMR (600 MHz, CDCl_3_) δ 7.63 (t, *J* = 1.6 Hz, 1H), 7.61 (t, *J* = 1.5 Hz, 1H), 7.33 (t, *J* = 1.7 Hz, 1H), 6.34 (br s, 2H), 4.25 (t, *J* = 7.1 Hz, 2H), 2.75 – 2.68 (m, 1H), 2.56 (t, *J* = 6.8 Hz, 2H), 1.94 (p, *J* = 6.8 Hz, 2H), 1.04 (d, *J* = 6.3 Hz, 6H); ^13^C NMR (150 MHz, CDCl_3_) δ 159.3 (d, *J* = 212.0 Hz), 156.5 (d, *J* = 20.3 Hz), 152.9 (d, *J* = 19.2 Hz), 143.6 (d, *J* = 2.7 Hz), 136.9, 136.5, 135.8, 134.6, 129.3, 118.4 (d, *J* = 3.7 Hz), 94.5, 48.8, 43.8, 42.1,30.2, 22.8; MS m/z 520.9 [M + H]^+^.

### COMPUTATIONAL METHODS

#### System preparation

The starting structure for all simulations of the glycosylated models of full-length GRP94, either with ATP or ligands bound at the active site, is crystal structure 5ULS.pdb^43^; both protomers were reconstructed without gaps by modeling the missing fragments with Modeller ^73^. As the original structure contains AMPPNP, to reconstruct ATP, the Nβ atom of the adenylyl-imidodiphosphate molecule that is present in each protomer’s active site was replaced by oxygen. To predict ligand binding modes and concomitant structural changes in the receptor for the designed inhibitors, the adenylyl-imidodiphosphate molecule was removed and ligands PU-WS12 and PU-WS13 were docked into the binding site using the Glide docking program (Glide, version 6.9, Schrödinger, LLC, New York, NY, 2021) using the Induced Fit Docking (IFD) protocol, based on Glide and the Refinement module in Prime. Docking calculations were performed in standard precision mode (SP) with the OPLS_2005 force field. No modifications were applied to the default settings. The best poses according to the docking score function were selected as a starting point for MD simulations. All hydrogens were added (or replaced post-docking) using AmberTools’ tleap utility (version 19) ^74^, with residues calculated to be in their standard protonation states at physiological pH, (as predicted by PROPKA, v. 3.1. ^75^). Glycans were connected to residues N62 and N217 using the glycoprotein construction tool http://www.glycam.org: The following sequence was used to build the sugar chain: DManp *α*1,2 DManp *α*1,6 [DManp *α*1,2 DManp *α*1,3] DManp *α*1,6 [DManp *α*1,2 DManp *α*1,2 DManp *α*1,3] DManp ß1,4 DGlcpNAc ß1,4 DGlcpNAc - Asn.

#### Forcefield parameters for molecular dynamics simulations and parametrization of PU-WS12 and PU-WS13

During Molecular Dynamics simulations (MD), protein residues are modeled using ff14SB forcefield ^76^ parameters, except glycans and glycosylated asparagines N62 and N217 which were modeled using the GLYCAM_06j force field ^77^. ATP is treated using *ad hoc* parameters by Meagher and coworkers^78^. Compounds PU-WS12 and PU-WS13 parametrization were carried out with the aid of AmberTools’ antechamber and parmchk2 ^74^. Derivation of point charges was carried out using the Gaussian16 program (www.gaussian.com) in conjunction with antechamber: first, compounds were structurally optimized at the B3LYP/6-31G(d) level of density functional theory (DFT); subsequently, ESP charges ^79^ around each atom are calculated based on the electrostatic potential, calculated at the Hartree-Fock/6-31G(d) level, and sampled over 10 shells per atom at a density of 17 grid points per square Bohr; final atomic point charges are assigned after RESP ^80^ fitting performed by antechamber. The chosen water model is TIP3P ^81^ which is compatible with parameters by Joung and Cheatham ^82^ chosen to treat Na^+^ cations used to neutralize the charge of the system.

#### Molecular dynamics simulations

MD simulations were carried out using the Amber suite (version 18) ^74^ with the Sander MD engine employed in the early preproduction stages (minimization, heating and equilibration) of each MD run, and the GPU-accelerated pmemd.cuda utility ^83^ thereafter. Three independent MD replicas (atomic velocities assigned from different random seeds) are carried out for each system, comprising minimization, heating, equilibration, and production. First, the prepared solute was energy-minimized in vacuo using 200 steps of steepest descent and an additional 200 steps using conjugate gradients. A cutoff of 10 Å was used for Coulomb and van der Waals interactions. The minimized system was then solvated using the TIP3P water model in a truncated octahedral box and rendered electroneutral by adding sodium counterions. To remove any bad contacts between solute and solvent, the system was minimized with a two-stage approach. In the first stage we restrained the solute coordinates using a 500 kcal mol^-1^Å^-2^harmonic constant force and minimized the positions of water and ions with 500 steps of steepest descent followed by 500 steps of conjugate gradient, a. Then, in the second stage, the entire system was minimized with 1000 steps of steepest descent and 1500 conjugate gradient steps without restraints. The temperature of the system was then increased from 0 to 300 K in the NVT ensemble, running a rapid heating step of 20 ps of MD with weak positional restraints on the solute (force constant of 10 kcal kcal mol^-1^Å^-2^) and the Langevin thermostat^84^ (collision frequency of 1.0 ps-1) to avoid any large fluctuations. All bonds involving hydrogen were constrained using the SHAKE^85^ algorithm. The systems were then equilibrated with no restraints for 500 ps with a 2 fs time step, with the SHAKE ^85^ algorithm and periodic boundary conditions in the NPT ensemble. Initial velocities for each replicate were obtained from a Maxwellian distribution at the initial temperature of 300 K. The temperature was kept constant with the Langevin thermostat (collision frequency of 1.0 ps-1) ^84^, and a constant pressure of 1 bar was introduced via Berendsen’s barostat (2 ps relaxation time) ^86^. The electrostatic interactions were treated using the particle mesh Ewald method^87^ with a cutoff of 10 Å. The same cutoff was used even for short-range Lennard-Jones interactions. Each production run (3 for each system) was extended to 1 μs with an identical setup to the final equilibration condition. Postprocessing and analysis of MD trajectories were conducted with the CPPTRAJ tool^88^ whereas the VMD platform was used for visual inspection^89^.

#### Analysis of MD trajectories

##### Distance fluctuations (DF) analysis

The replicates for each system were combined into 1 comprehensive metatrajectory (one metatrajectory for each system reported in **Table 1**). The matrix of the fluctuations of pairwise amino acid distances has been obtained by calculating the DF parameter for any pair of residues ^44^. This parameter is defined as *DF_ij_* = (*d_ij_* – 〈*d_ij_*〉))^2^, where *d_ij_* is the (time-dependent) distance of the Cα atoms of amino acids i and j, and the brackets indicate the time average over the trajectory. The DF matrix was calculated for the each of the meta-trajectory of all the simulated systems.

##### Analysis of contacts

Analysis of contacts between the glycans (or the ligands) and the protein over the trajectories have been performed using the *nativecontacts* command implemented in the *CPPTRAJ* program distributed within the *AmberTools* suite (version 19) ^74^. To count as a contact, the distance between residues i and j in a structure should be lower than 7 Å. The frequency of each contact has been determined with in-house python script. Calculations were carried out independently for each sugar chain (or for each ligand).

##### Root-mean-square-deviation (RMSD)

RMSD of protein backbone atoms was computed using the *rmsd* command implemented in the CPPTRAJ program distributed within the *AmberTools* suite (version 19) ^74^. Alignment of each snapshot of the analyzed trajectory using the coordinates of the corresponding backbone atoms of the crystal structure (PDB code 5ULS,^43^ as a reference was done before RMSD calculations. Calculations was carried out independently for each protomer.

##### Cluster analysis

Cluster analysis was carried out using hierarchical agglomerative algorithm from *cluster* command in CPPTRAJ distributed within the *AmberTools* suite (version 19) ^74^ Clustering was carried out in the following way: first, all snapshots from the metatrajectories of each simulated system are aligned using the coordinate of backbone atoms of the entire protein of the starting conformation, then the coordinated of the backbone atoms of the lid in each snapshot are used to define the different conformational families, using as distance metric option the RMSD with the *nofit*option. For each simulated system, 10 representative conformations were collected, with the *complete-linkage* method, which uses the maximum distance between members of two clusters.

##### Docking calculations

Compound Bnlm (**Figure 1**) was docked with the GLIDE docking program (Glide, version 9.6, Schrödinger, LLC, New York, NY, 2022) at the ATP binding site of the central structures of the most populated conformational clusters obtained from the trajectories of the fully glycosylated and the WT forms bound to ATP, respectively 2GlycATP and NoGlycATP. Rigid receptor and flexible ligand docking calculations were performed in standard precision mode (SP) with the OPLS_2005 force field, nonplanar conformations of amide bonds were penalized, van der Waals radii were scaled by 0.80, and the partial charge cut-off was fixed to 0.15. No further modifications were applied to the default settings.

### QUANTIFICATION AND STATISTICAL ANALYSIS

Any quantification and statistical analysis that were applied in the experiments are described in Method details, as well as in the main and supplemental figure legends.

## Notes

### Competing Interest Statement

The authors have declared no competing interest.

