## Supplemental Figures and Tables for "How Aberrant *N*-Glycosylation Can Alter Protein Functionality and Ligand Binding: an Atomistic View"

### Supplementary Figure 1

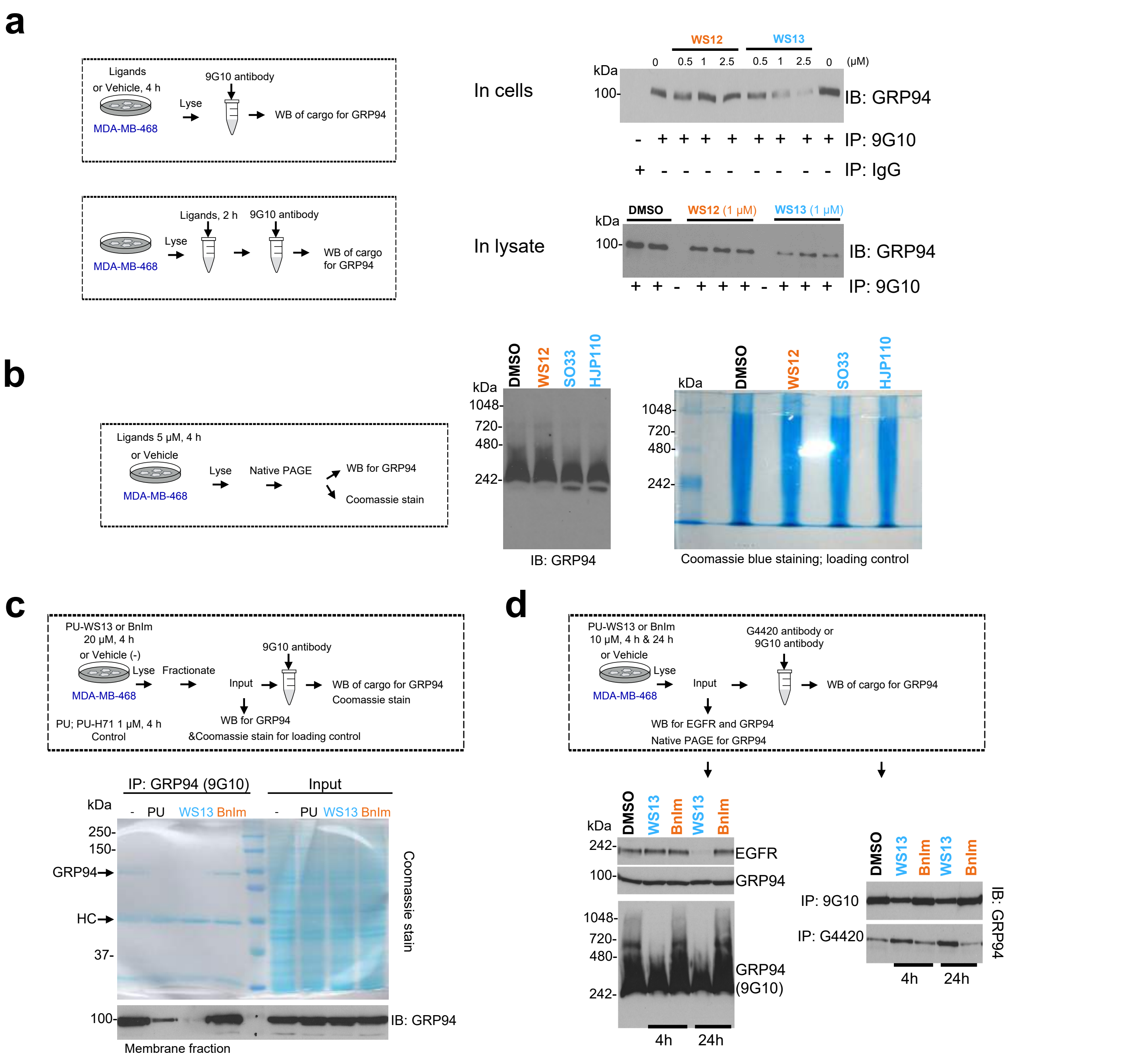


**Figure S1. Testing paradigm for small molecule activity via $^{Glyc62}$GRP94, Related to Figure 1.**
(a) Immuno-capture of GRP94 was performed as shown in the schematic. MDA-MB-468 cells or cell homogenates were treated with ligands or vehicle (DMSO) as indicated. Top panel, for 4 h at 0.5, 1 and 2.5 μM and lower panel, for 2 h at 1 μM, prior to immunocapture and WB of the cargo. Experiments were repeated thrice with similar results. IgG, isogenic control.
(b) Native-PAGE followed by immunoblotting for GRP94 (left) or Coomassie staining (right). MDA-MB-468 cancer cells were treated with vehicle (DMSO) or indicated ligands (at 5 μM for 4 h) prior to lysing and analysis. Gels are representative of three independent results.
(c) As in (a) for MDA-MB-468 cells treated with ligands (at 20 μM for 4 h) or vehicle (DMSO) prior to immunocapture and WB of the cargo. The membrane fraction was used for immunocapture. The WB and Coomassie stained gels of the input are shown as control for equal protein amount, as used for immunocapture. Gels are representative of three independent results.
(d) Functional and biochemical analysis of PU-WS13 and BnIm in a battery of assays that test productive $^{Glyc62}$GRP94 engagement in cells. MDA-MB-468 cells were treated for 4h or 24 h with inhibitors (at 10 μM) prior to analysis, as indicated in the schematic. $^{Glyc62}$GRP94 inhibits EGFR internalization and degradation, and $^{Glyc62}$GRP94 inhibition suppresses this effect (see EGFR steady-state level analysis by WB). GRP94 conformational modulation by ligands was detected by immunocapture and Native PAGE, as indicated.

### Supplementary Figure 2

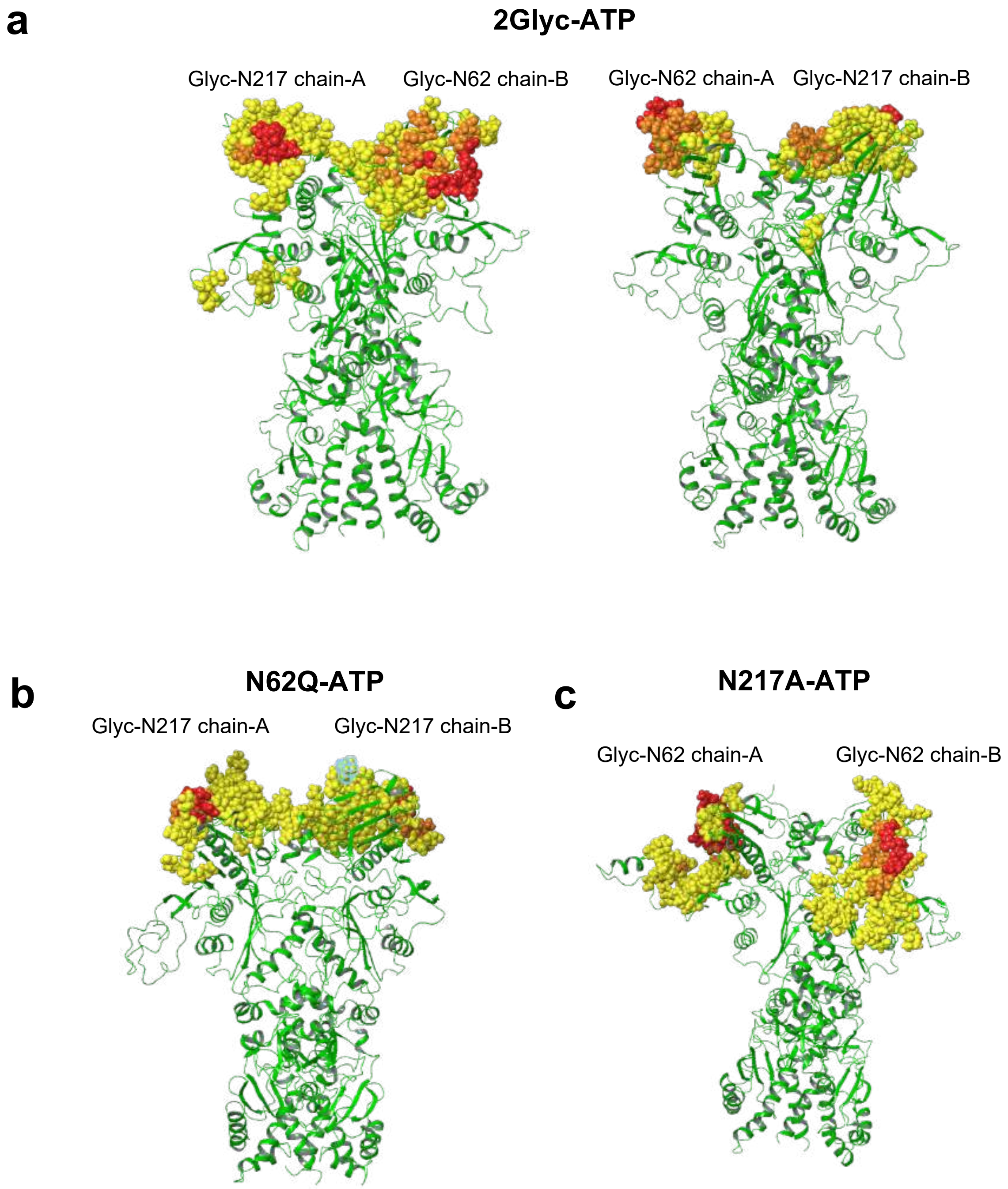


**Figure S2. Binding of ATP to the GRP94 variants, Related to Figures 2 and 3.**
(a) Contact between sugar chain and protein for 2GlycATP.
(b) Contact between sugar chain and protein for N62Q-ATP.
(c) Contact between sugar chain and protein for N217A-ATP.
In yellow we represent residues of the proteins that make contact with the sugars 20% to 50% of the time, orange 50% to 70%, red 70% to 100%.
For Supplementary Tables see Supplementary data provided electronically (Data_Glyc_Contacts.zip).

### Supplementary Figure 3

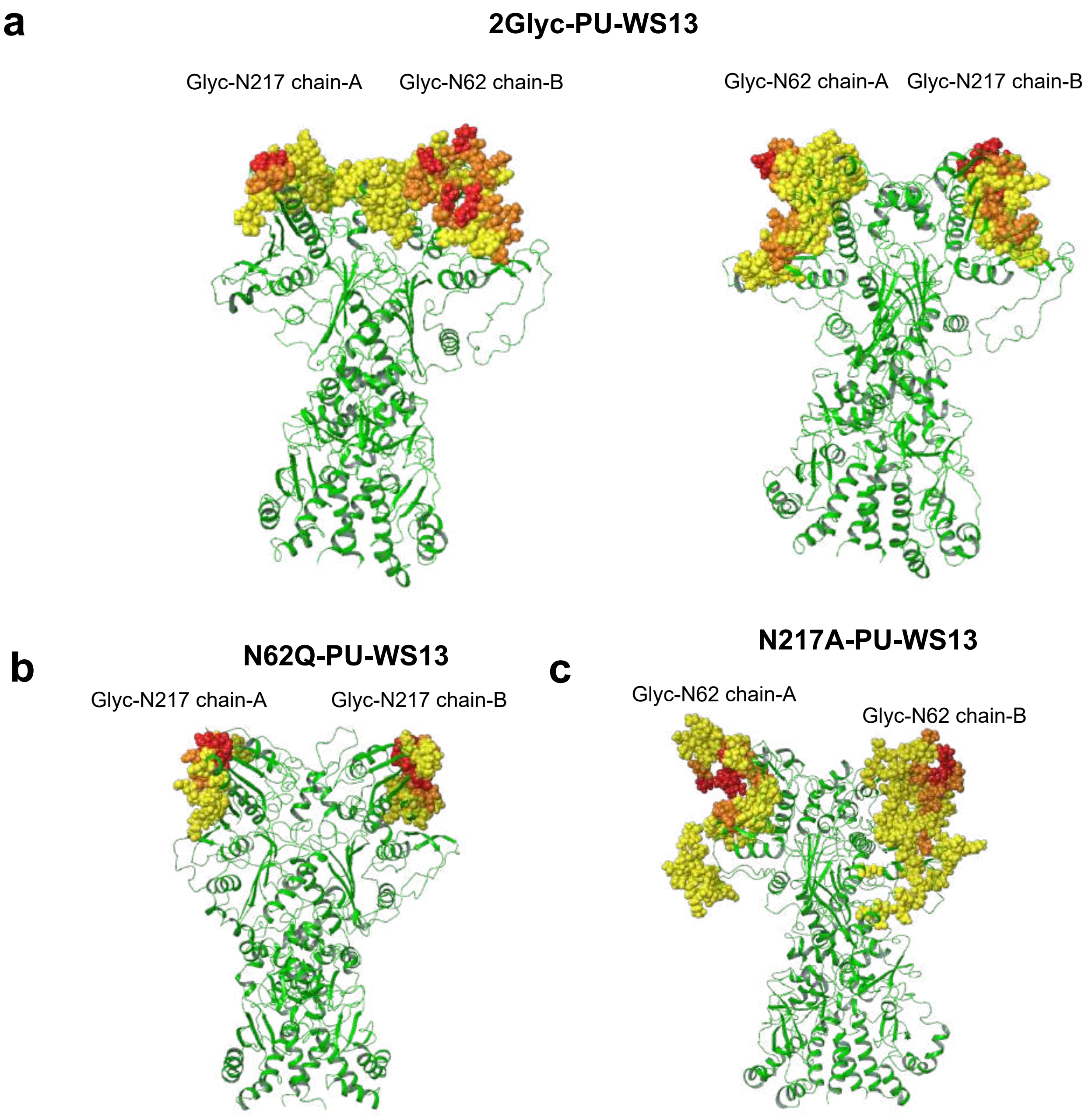


**Figure S3. Binding of PU-WS13 to the GRP94 variants, Related to Figures 2 and 3.**
(a) Contact between sugar chain and protein for 2Glyc-PU-WS13.
(b) Contact between sugar chain and protein for N62Q-PU-WS13.
(c) Contact between sugar chain and protein for N217A-PU-WS13.
In yellow we represent residues of the proteins that make contact with the sugars 20% to 50% of the time, orange 50% to 70%, red 70% to 100%.
For Supplementary Tables see Supplementary data provided electronically (Data_Glyc_Contacts.zip).

### Supplementary Figure 4

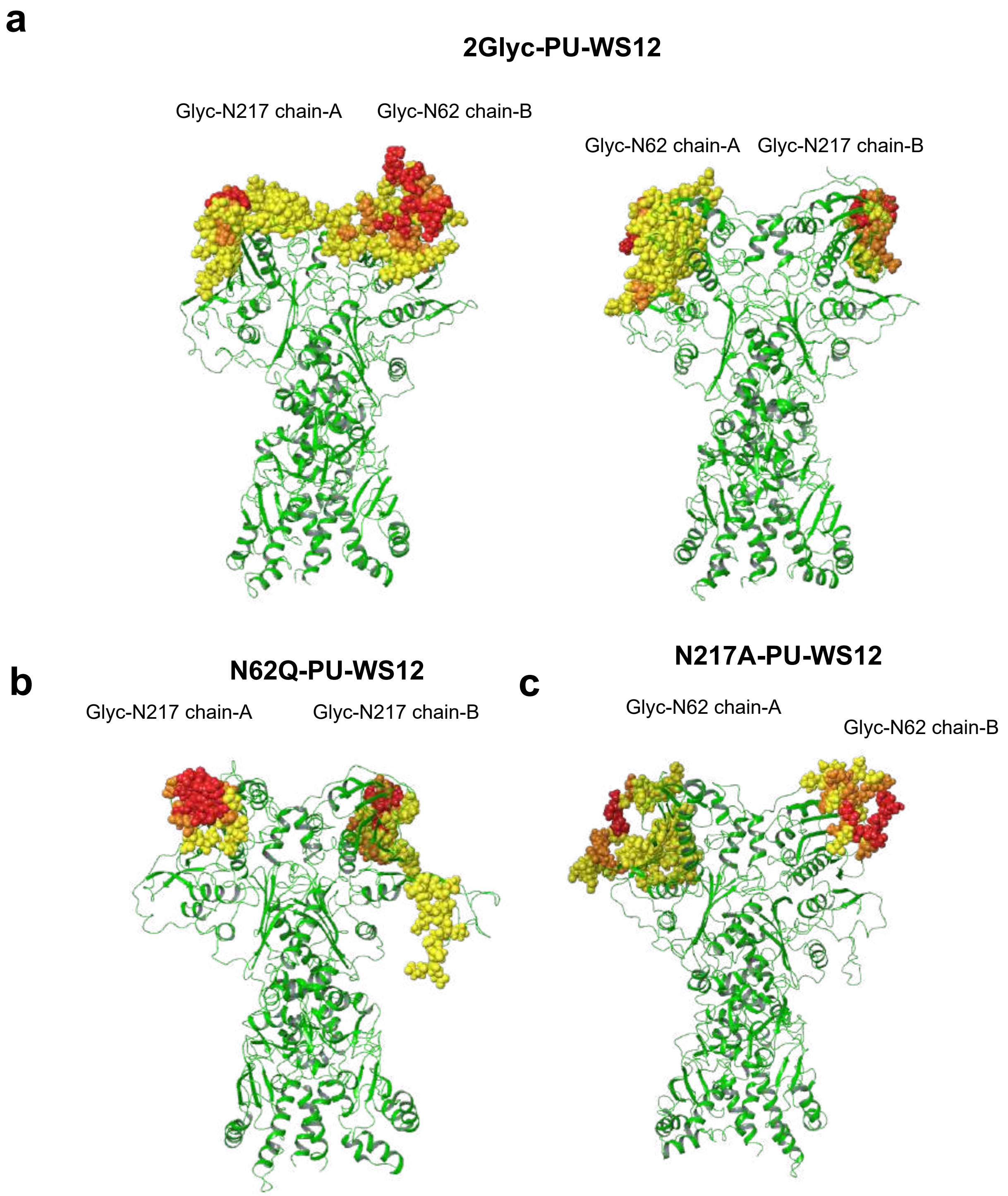


**Figure S4. Binding of PU-WS12 to the GRP94 variants, Related to Figures 2and 3.**
(a) Contact between sugar chain and protein for 2Glyc-PU-WS12.
(b) Contact between sugar chain and protein for N62Q-PU-WS12.
(c) Contact between sugar chain and protein for N217A-PU-WS12.
In yellow we represent residues of the proteins that make contact with the sugars 20% to 50% of the time, orange 50% to 70%, red 70% to 100%.
For Supplementary Tables see Supplementary data provided electronically (Data_Glyc_Contacts.zip).

### Supplementary Figure 5

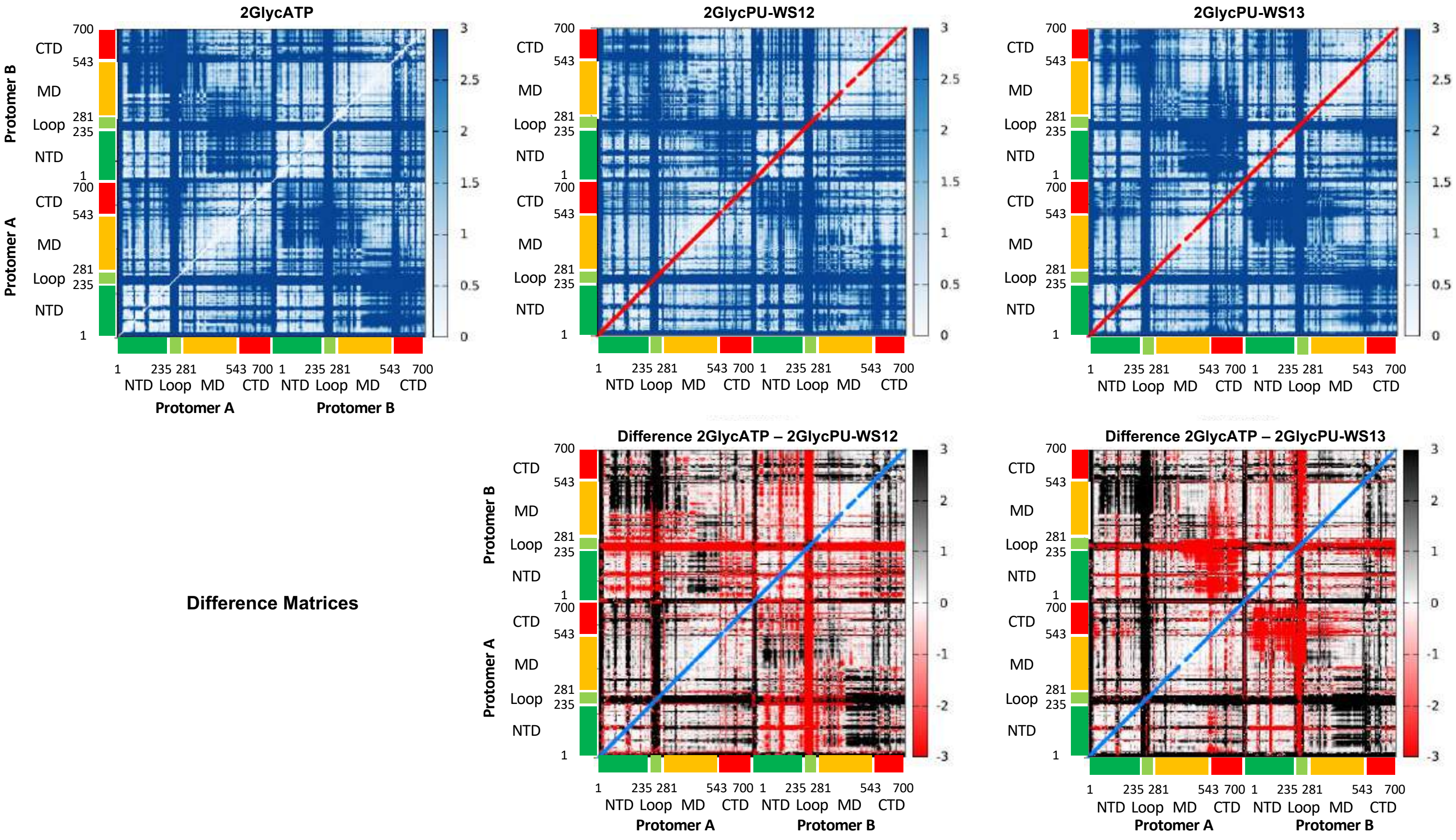


**Figure S5. Distance Fluctuation (DF) matrix for the 2Glyc state, bound to ATP, PU-WS12 or PU-WS13, Related to Figures 5 and 6.**

The reference DF matrix for the fully glycosylated ATP-state of GRP94 and the difference-matrices obtained by subtracting the DF values of another state (different ligand, mutant, or mutant+ligand) from this reference, are presented. This type of representation is intended to highlight the regions where the different chemical perturbations determine the largest impact on the internal dynamics of the protein: the color code indicates lower coordination (higher dynamic flexibility, less ordered motions, easier disruption of local interaction networks) with respect to the reference state in red, and higher coordination (increased communication between different areas) in black. Matrix representation of the residue-pair DFs was calculated from the various trajectories. In each subpanel, the top row represents the original matrix, while the bottom row represents the difference matrix obtained subtracting the values for a certain mutant/ligand state from the reference matrix corresponding to the 2GlycATP state. If residue-pair fluctuations are larger (lower coordination, larger flexibility) in the specific mutant/ligand state than in the reference state, the corresponding value in the difference is depicted in red; if residue-pair fluctuations are smaller (higher coordination, lower flexibility) in the specific mutant/ligand state than in the reference state, the corresponding value in the difference is depicted in black.

### Supplementary Figure 6

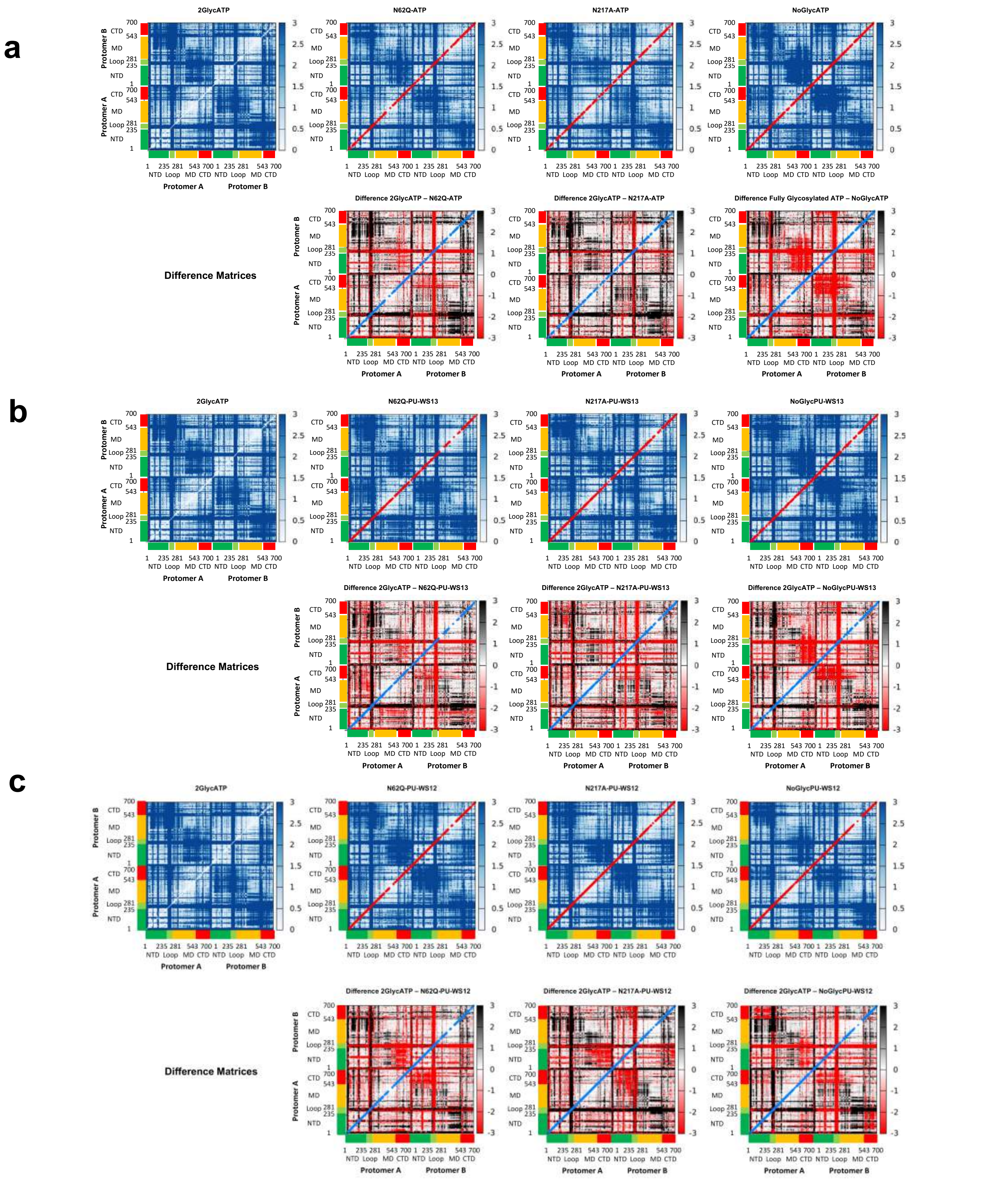


**Figure S6. Distance Fluctuation matrix for different mutants, including the unglycosylated Wild Type, in complex with various ligands, Related to Figures 5 and 6.**

(a) DF matrix for GRP94 variants in complex with ATP.

(b) DF matrix for GRP94 variants in complex with PU-WS13.

(c) DF matrix for GRP94 variants in complex with PU-WS12.

### Supplementary Figure 7

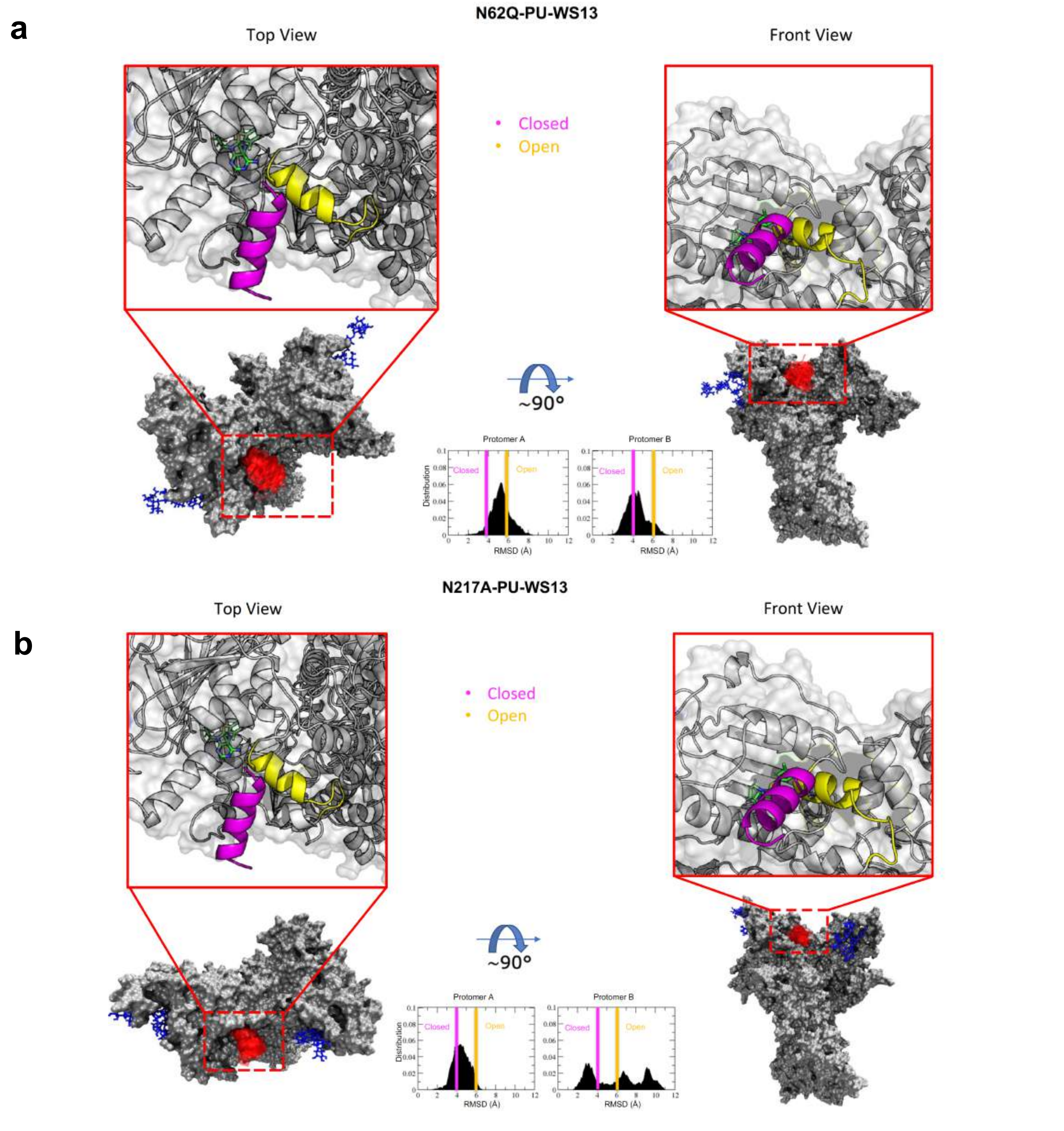


**Figure S7. Conformational dynamics of the ATP-lid covering the active site of GRP94 variants and their interactions with sugar chains in the open or closed forms, Related to Figures 7, 8 and 9.**

(a) Analyses for the N62Q-PU-WS13 system.
(b) Analyses for the N217A-PU-WS13 system.
Two orientations are depicted. ATP-lid (at several times, 300 frames) and glycans are represented with red and blue lines, respectively. The insets show the distributions of RMSD values from the conformation presented in crystal structure PDB 5ULS.pdb. The blue and yellow vertical lines show the RMSD threshold for the closed and open states, respectively.

### Supplementary Figure 8

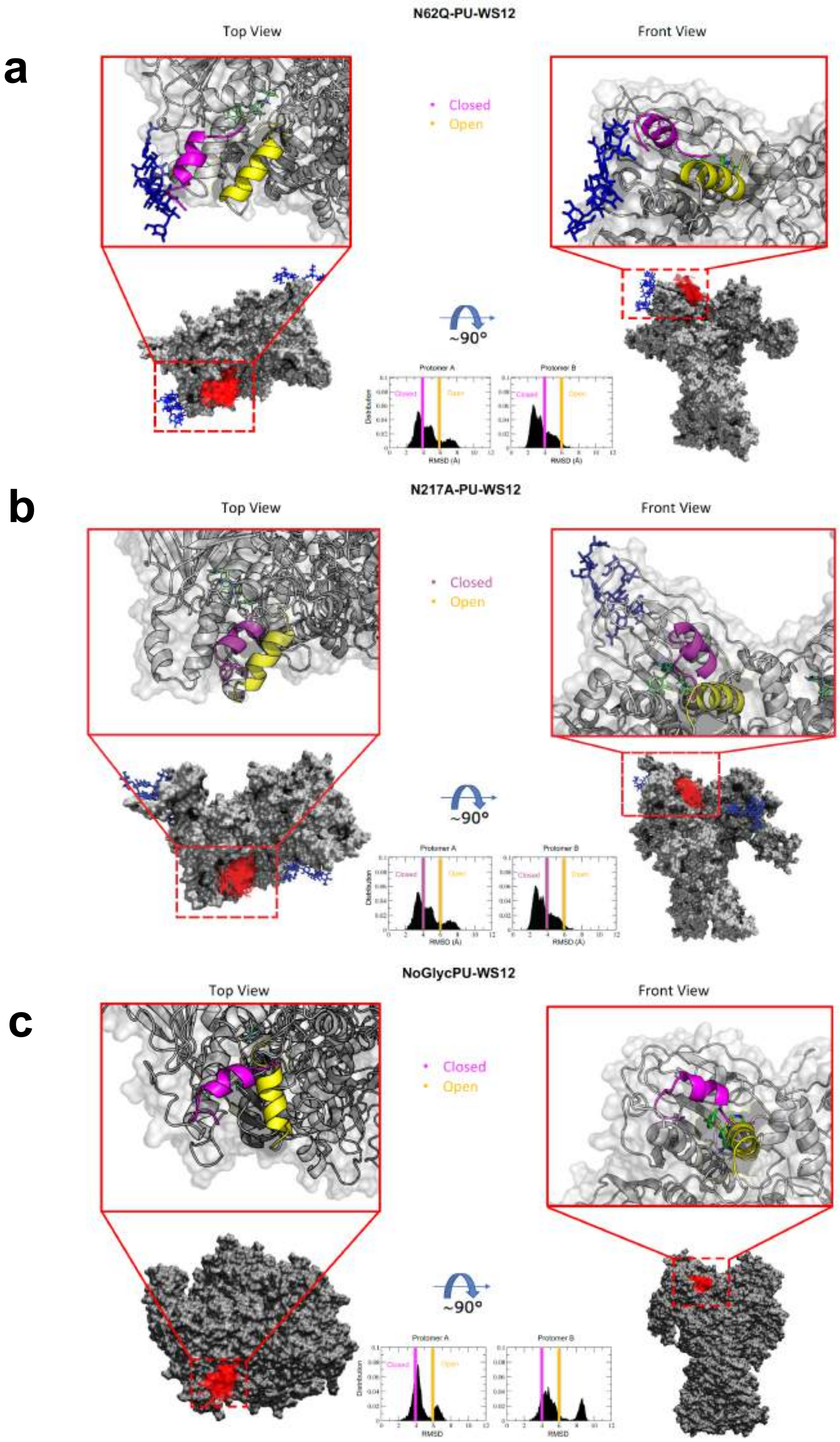


**Figure S8. Conformational dynamics of the ATP-lid covering the active site of GRP94 variants and their interactions with sugar chains in the open or closed forms, Related to Figures 7, 8 and 9.**

(a) Analyses for the N62Q-PU-WS12 system.
(b) Analyses for the N217A-PU-WS12 system.
(c) Analyses for the NoGlyc-PU-WS12 system.

Two orientations are depicted. ATP-lid (at several times, 300 frames) and glycans are represented with red and blue lines, respectively. The insets show the distributions of RMSD values from the conformation presented in crystal structure PDB 5ULS.pdb. The blue and yellow vertical lines show the RMSD threshold for the closed and open states, respectively.

### Supplementary Figure 9

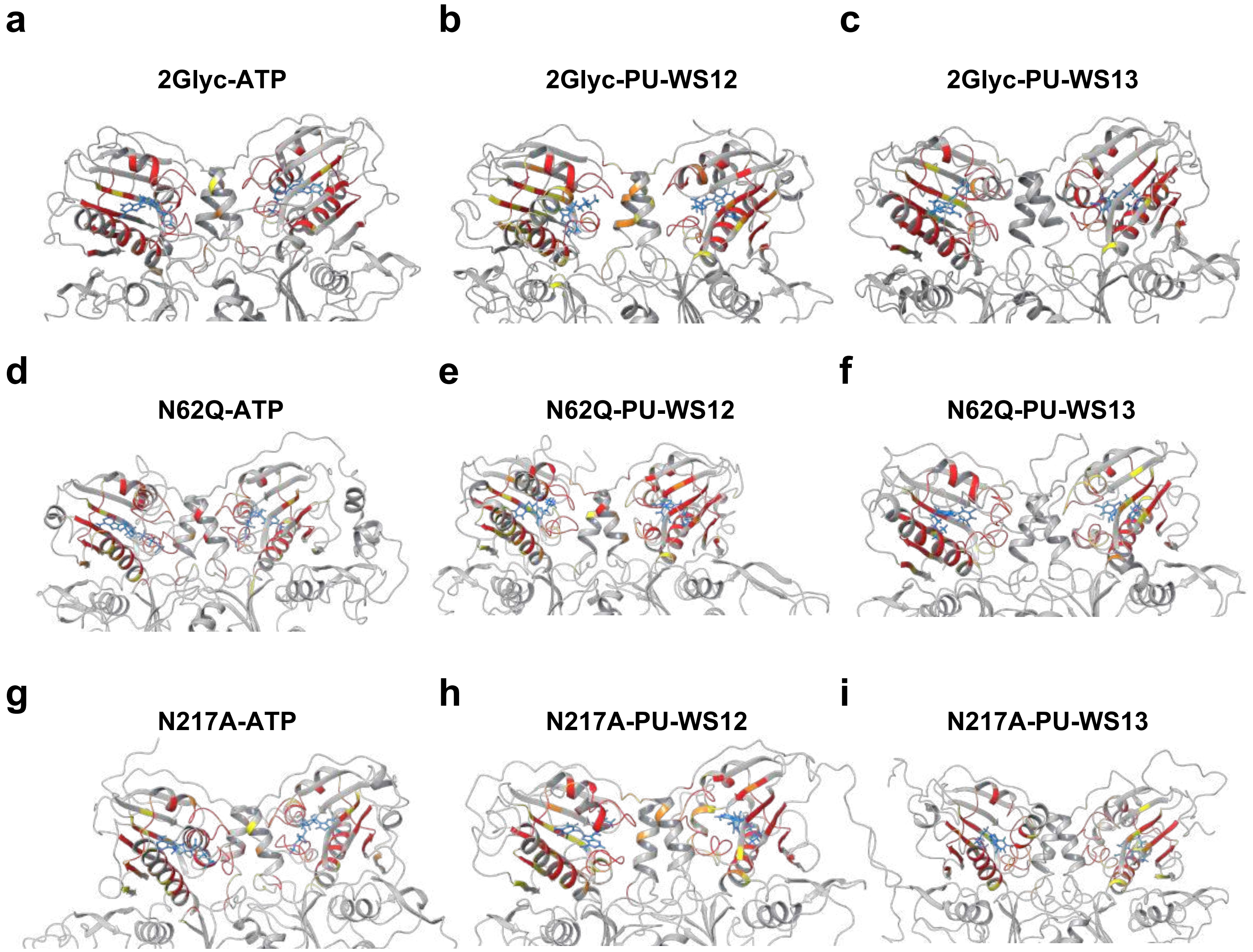


**Figure S9. Contact between ligand and surrounding residues of the protein for GRP94 variants in complex with ligands.**

(a) The 2Glyc-ATP system.
(b) The 2Glyc-PU-WS12 system.
(c) The 2Glyc-PU-WS13 system.
(d) The N62Q-ATP system.
(e) The N62Q-PU-WS12 system.
(f) The N62Q-PU-WS13 system.
(g) The N217A-ATP system.
(h) The N217A-PU-WS12 system.
(i) The N217A-PU-WS13 system.

In yellow we represent residues of the proteins that make contact with the ligand 20% to 50% of the time, orange 50% to 70%, red 70% to 100%.

For Supplementary Tables see Supplementary data provided electronically (Data_Ligands_Contacts.zip).

### Supplementary Figure 10

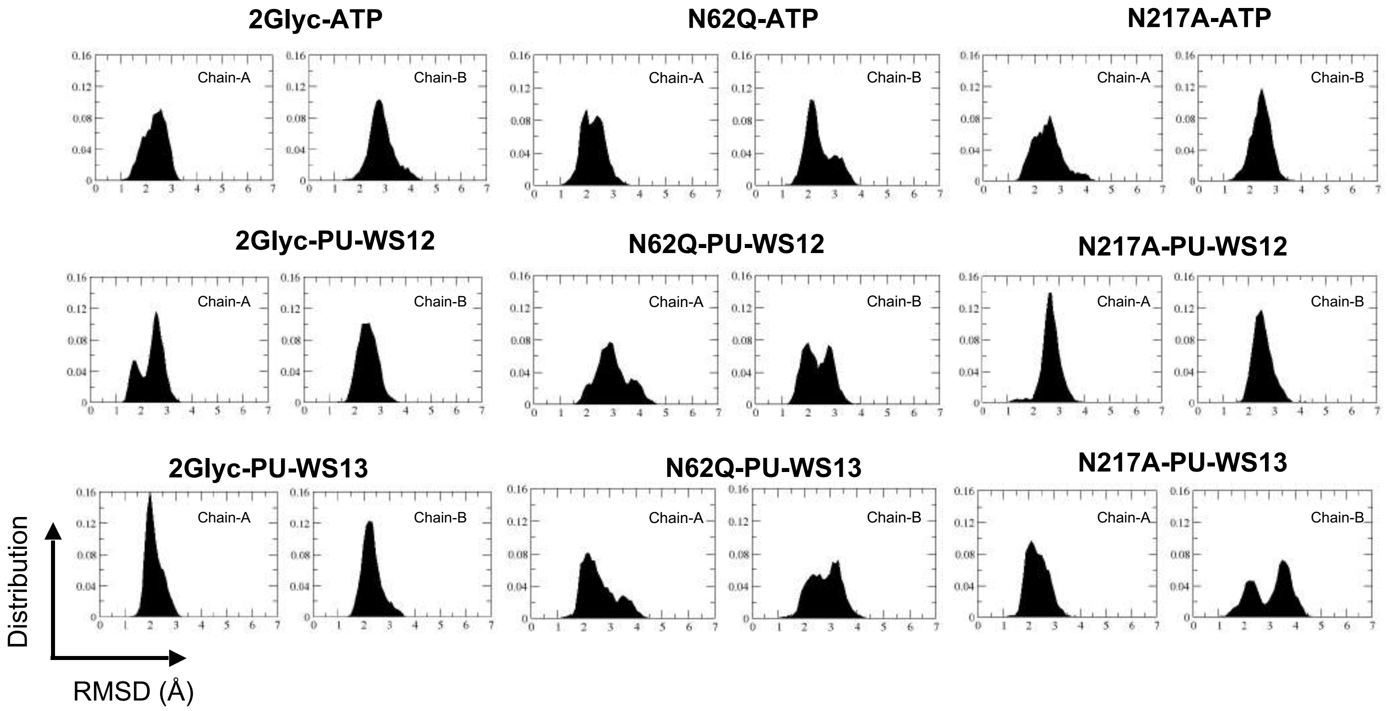


**Figure S10. RMSD distribution over the entire metatrajectory for indicated GRP94 variant-ligand systems.**

Calculation was carried out independently for each protomer and focused on the residues making up the active site compared to their respective positions in the ATP-bound crystal structure of GRP94 (PDB: 5ULS).

### Supplementary Figure 11

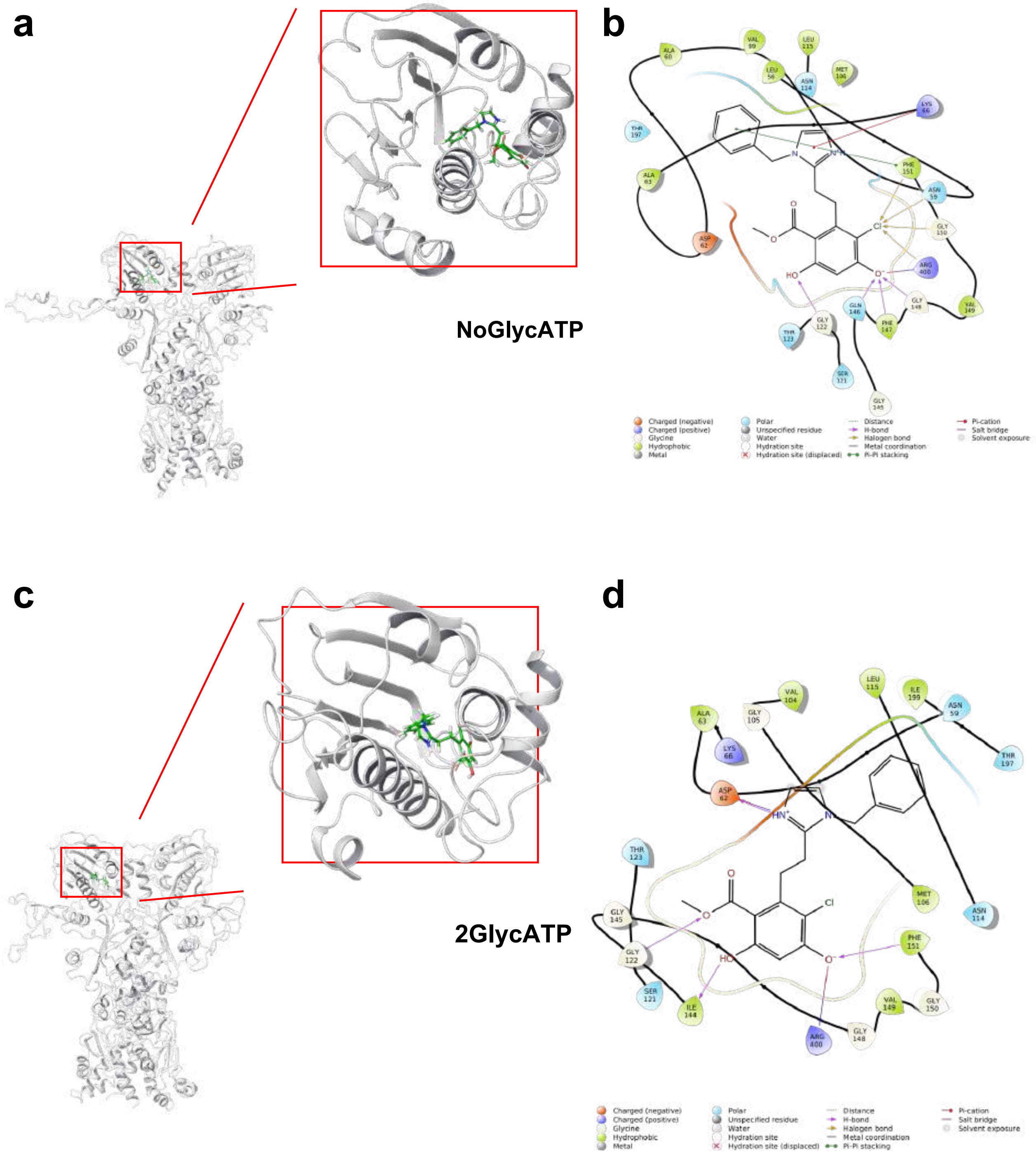


**Figure S11. BnIm docking analysis in GRP94 variants.**
(a) Representative structure for the best scoring docking pose of BnIm into the active site of GRP94 NoGlycATP. Zoom on NTD.
(b) 2D-interaction di agram based on the best-scoring docking pose of BnIm as in (a). Figure generated with Schrodinger's Maestro suite.
(c) Representative structure for the best scoring docking pose of BnIm into the active site of GRP94 2GlhycATP. Zoom on NTD.
(d) 2D-interaction diagram based on the best-scoring docking pose of BnIm as in (c).
See Supplementary data provided electronically (Data_Docking_BnIm.xlsx)
